# Engineering Regionally-Activated Drugs for Neuroscience

**DOI:** 10.1101/2023.10.12.562069

**Authors:** Zhimin Huang, Andrei Mitrofan, Shirin Nouraein, Clark Horak, Joon Pyung Seo, Manwal Harb, Rongshu Jin, Jerzy O. Szablowski

## Abstract

The brain is comprised of multiple regions performing distinct functions. Within each of these regions, there are multiple cell types that can affect brain physiology. Finally, within each cell there are multiple signaling pathways, that, when activated or inhibited, control the cell’s activity, and consequently the brain function. For these reasons, methods that can control the brain with regional, cell-type, and molecular precision have been widely used in neuroscience. However, so far, achieving sustained control over a brain region with that level of specificity relied either on gene delivery or placement of invasive devices. While gene therapy holds great promise, the risks of genomic integration, vector toxicity, vector-directed immune response, high cost, and gene delivery to the brain pose significant challenges. On the other hand, invasive devices enable site-specific delivery of drugs but can also surgically damage the modulated brain region, carrying risks of infection and hemorrhage. Here, we present a new approach that can provide multi-day, noninvasive, site-specific control over specific cell types in the brain without the need to use invasive devices or gene delivery. To achieve this, we introduce a new paradigm called Regionally Activated Interstitial Drugs, or **RAID**, which delivers a protein-based catalytic centers, or RAID enzymes, to the brain using focused ultrasound blood-brain barrier opening. This catalytic center is designed to attach to the interstitial space in the brain where it remains for days after initial delivery. While the catalytic center is present in the brain, it can locally process an inert BBB permeable prodrug into an active drug, resulting in localized therapy. Our proof-of concept studies demonstrated that the engineered RAID enzymes can retain activity in the brain parenchyma for several days, allowing for noninvasive site-specific induction of neuronal activity that was sufficiently potent to elicit behavioral effects. Overall, the RAID paradigm enabled noninvasive, tunable, temporally-resolved, site-specific, non-genetic, neuromodulation over multiple days. The RAID paradigm is versatile and can be applied to any enzyme and prodrug pair to control various aspects of central nervous system physiology.

## INTRODUCTION

The most common strategy to control brain activity is to use small molecule drugs targeted to specific cellular receptor. However, the brain consists of regions that can be distinguished by their anatomical features, functions^1^, and disease involvement^2-5^. Small molecule drugs diffuse throughout the brain and act without spatial precision resulting in off-target effects^6-11^. On the other hand, controlling specific brain sites with molecular precision enabled critical advances in neuroscience and allowed for control of specific behaviors^12-14^. If these approaches became feasible for clinical treatments, they would revolutionize how we treat neurological and psychiatric disorders. However, the majority of these spatially-specific neuromodulation tools rely on surgical delivery of molecules^15,16^, or device implantation^17-19^, which are invasive and damage the tissue, or on gene therapy, which is expensive and typically restricted to a single administration due to immune response directed to the most commonly used gene delivery vectors^20-22^. Finally, when neuromodulation is powered with a device, these devices can be bulky, restrict the motion of research animals, are infeasible when targeting large swathes of the brain, and in the case of clinical translation – can present social challenges. For example, optogenetics can control neurons^12^ with tunable levels of activation, for extended periods of time, and with spatial, cell-type, and temporal precision. However, optogenetics typically requires gene delivery, an externally-powered device, and the implantation of an optical fiber in the proximity of the stimulated site due to the limited penetration of light through the tissue^18,19,23^. When optogenetics is used in large brain regions, it requires implantation of a large numbers of such fibers^24-26^ leading to tissue damage. Noninvasive long-term neuromodulation with spatial, cell-type, and temporal precision, and without an external wearable device is possible with Acoustically Targeted Chemogenetics (ATAC)^27^. However, ATAC still requires gene delivery to the brain with all its potential limitations and challenges. In ATAC, adeno-associated viral vectors (AAVs) are delivered systemically and then into specific brain regions with the use of focused ultrasound bloodbrain barrier opening (FUS-BBBO). These vectors encode chemogenetic receptors^13,28^ which can then control the transduced neurons in response to a systemically administered drug. These approaches can have significant impact, however, in many cases there is a need to confirm the utility of targeting the specific brain site, and controlled molecular pathway without invasive devices, or expensive ‘single-use’ gene therapy.

A partial resolution of this need is a systemic delivery of therapeutics combined with FUS-BBBO^29-32^. It allowed for non-invasive, tunable, spatially-, cell-type, and temporally defined brain stimulation. However, it only allowed such neuromodulation for a limited amount of time while the BBB stayed open. In this approach, BBB-impermeable small molecule drugs are administered systemically and only enter the brain sites targeted by FUS-BBBO. While this approach leads to systemic effects, just as a typical drug administration, it can still be useful in determining the contribution of specific brain regions on behavior^33-35^. Unfortunately, the FUS-BBBO-delivered small-molecule drugs are typically retained in the brain for short periods of time ranging from minutes to hours^30,33,36,37^. In consequence, any neuromodulatory effects of drugs delivered with FUS-BBBO are short-lived and inherently confounded by the presence of disrupted BBB, which closes over 6-24 h^38-40^. Such short duration of action and reliance on the continuous presence of disrupted BBB limits practicality of this approach. For example, drugs for psychiatric disorders such as schizophrenia, or obsessive-compulsive disorder, are administered multiple times a day^41-43^. It would be impractical and expensive to perform daily noninvasive FUS-BBBO procedures, as they currently take several hours and require magnetic resonance imaging sessions for spatially precise targeting^44,45^, or use less spatially precise implantable devices^46^. In another example, confirmation of the seizure focus before surgical resection is a major challenge^47,48^. Such resection often leads to side effects, while having limited effects on the seizure frequency and severity^49,50^. Because seizures occur randomly and infrequently, the patients are typically observed in the epilepsy unit for several days at a time, indicating a need for site-specific silencing of the seizure foci over that period of time^51,52^. If control of specific brain regions using non-genetic, non-invasive, spatially precise methods was possible, it would enable site-specific therapy of the circuits affected in psychiatric or neurological disorders, and would enable either treatment or confirmation of effects of shutdown of specific seizure foci before surgical resection, device implantation, or other more involved therapies.

Unfortunately, such methods have not yet been found. In theory, delivery of nanoparticles loaded with drugs using FUS-BBBO could extend the timeline of drug release, but the nanoparticles tested so far are too large to be efficiently delivered with FUS-BBBO at safe ultrasound pressures^53-56^. Additionally, such an approach would lead to the release of drugs in a pre-determined pattern without on-demand tunability in response to a patient’s input or condition. Ability to tune the drug dose is common in treatment of psychiatric and neurological disorders^57,58^. In consequence, the FUS-BBBO-based drug delivery faces a conundrum - either accept short-term drug action or compromise the cost and safety to use long-acting therapies. Here, we propose a third way that achieves the best of both worlds, where we use a non-genetic approach that can be safely delivered to the brain with FUS-BBBO while also allowing for long-term drug action and tunability.

To achieve this, we demonstrate a new versatile concept for noninvasive, site-, molecular-, and temporally-specific neuromodulation that does not involve the challenges of genetically-encoded components or nanoparticles. This paradigm, called Regionally Activated Interstitial Drugs, or RAID, enables multi-day, tunable neuromodulation using molecules that are compatible with FUS-BBBO delivery **(Fig. 1a**). In RAID, we use FUS-BBBO to deliver an engineered enzyme to a localized brain region. That enzyme binds to the interstitial space within the brain, where it is retained for extended periods of time.

**Fig. 1:**
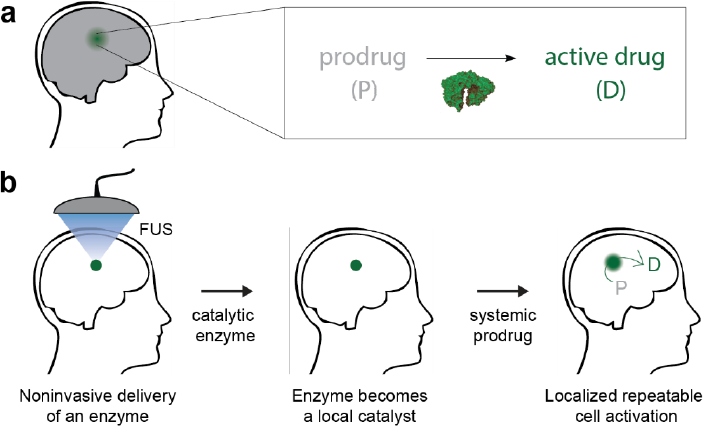
The concept of site-specific non-genetic therapeutics. **a)** An enzyme delivered to the selected site of brain specifically converts an inactive but BBB permeable prodrug (grey) into a neuroactive drug (green). We call this concept Regionally Activated Interstitial Drugs, or RAID. **b)** In RAID, catalytic enzymes are delivered to the brain via FUS-BBBO, where they bind within the brain interstitium to increase their residence time. While the enzyme is present, neuronal activity around the enzyme can be modulated by conversion of a systemically-supplied BBB-permeable prodrugs (P) into active drugs (D).

When present, this enzyme acts as a local drug factory - it converts an inert systemically-supplied and BBB-permeable prodrug into an active neuromodulatory drug (**Fig. 1b**). RAID is compatible with various drug doses and could be tunable on demand – by varying the dose of the prodrug, RAID could achieve stronger or weaker neuromodulation. In our studies, RAID enzymes stay within the brain and remain functional for at least several days. We also have shown that RAID can exert spatially-specific neuromodulation sufficient to control specific behaviors. Overall, RAID can be used to evaluate the effects of site-specific modulation of neuronal activity within the brain, while using a combination of human-derived and FDA-approved molecules. The RAID concept is versatile and can be applied to different prodrug-enzyme pairs, to control different signaling pathways and cell lines, in different brain regions that can be targeted with FUS-BBBO. Additionally, in our proof of concept, we used FDA approved molecules and a protein naturally present in humans, to showcase the potential utility of this concept in clinical research.

## RESULTS

### Retention of a native bioluminescent protein in the brain after FUS-BBBO delivery

To evaluate the retention of enzymatic activity in the brain, we delivered a model protein enzyme that is easily detectable *in vivo* and allows for facile tracking of the kinetics of its retention - RLuc8.6 luciferase. RLuc8.6 converts coelenterazine substrate into coelenteramide and in the process releases photons^59^ that can then be noninvasively detected in the brain with an intravital imaging system (IVIS) equipment (**Fig. 2a**). We injected the purified RLuc8.6 (150 mg/kg) intravenously and then used FUS-BBBO to deliver it deliver it unilaterally into the caudate putamen (CPu) of mice. We confirmed the spatially-specific delivery by immunostaining in brain sections (**Fig. 2b** and **Supplementary Fig. 1**). We found significantly higher immunostaining signal in the FUS target region (left striatum) at 1h post-delivery compared to the untargeted contralateral site (**Fig. 2c**, *n* = 4 mice, *P* = 0.0022, ratio paired t test (two-tailed)). To evaluate whether RLuc8.6 retained enzymatic activity in the brain interstitium, we measured the *in vivo* activity of RLuc8.6 at 4 different timepoints using IVIS. We found detectable RLuc8.6 enzymatic activity that was enhanced by FUS-BBBO in the brain after 1h (**Fig 2d**), and observed that opening the BBB in 1% of the brain volume led to 2.2(±0.3)-fold higher signal when averaged over the entire brain compared to RLuc8.6 alone without FUS (**Fig 2e**, *n* = 6 mice per group, *P* = 0.0074, one-way ANOVA test, with Tukey’s honestly significant difference (HSD) test). Considering the calculated volume of the single BBB opening site compared to the whole brain volume, the upper threshold of delivery should equal approximately 220-fold enzyme concentration increase within the targeted site compared to untargeted controls, which is consistent with previous studies of FUS-BBBO delivery for molecules of this size^60,61^. However, we also found that the IVIS signal decayed over time, and the enzyme levels were statistically indistinguishable from controls from 24 hours after FUS-BBBO onwards (**Supplementary Fig. 2**).

**Fig. 2:**
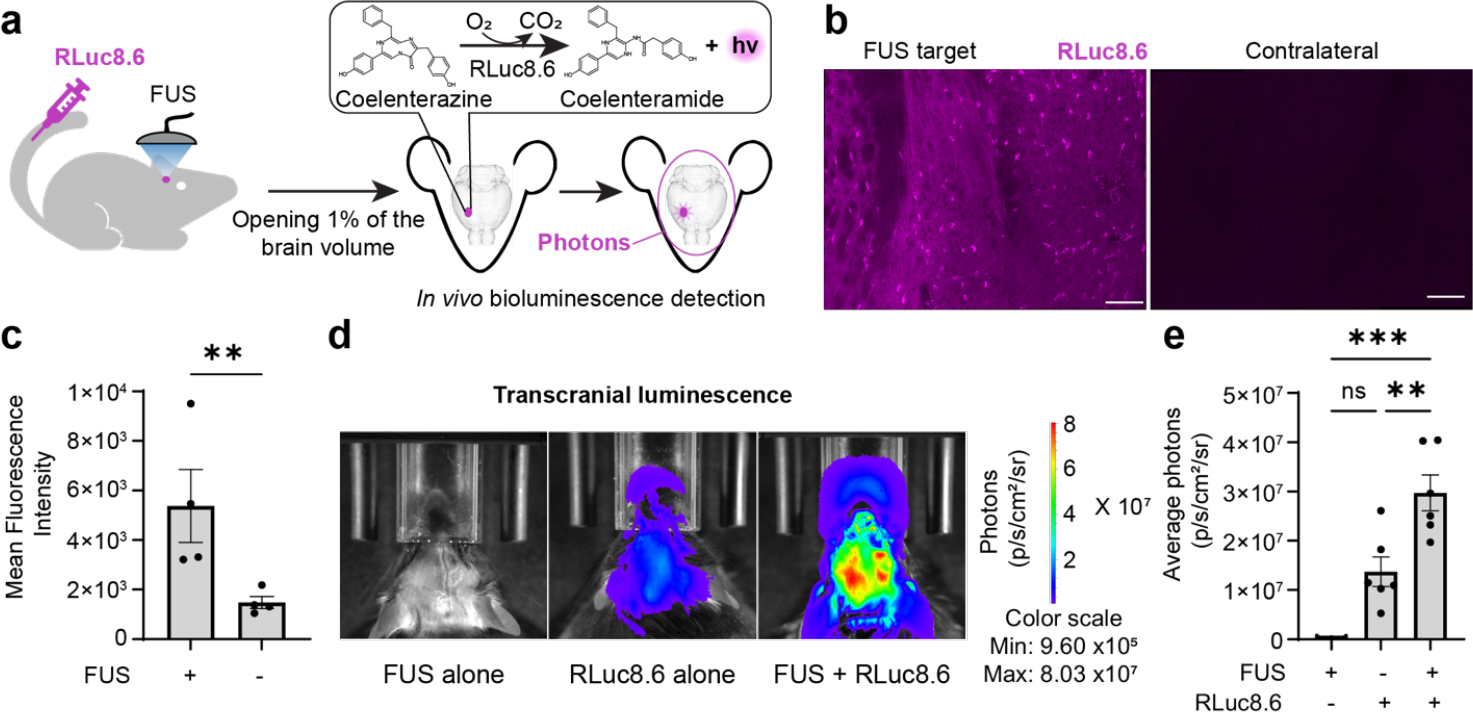
Noninvasively delivered enzyme retains activity in the brain interstitium. **a)** We delivered a recombinant luciferase, RLuc8.6, to mouse brain through FUS-BBBO and conducted bioluminescent imaging to assess the efficacy of targeted enzyme delivery and its activity. The activity was induced following an intraperitoneal injection of the Rluc8.6 substrate - coelenterazine (CTZ). **b)** Representative immunostaining images for RLuc8.6 (purple) in a mouse 1h after FUS-BBBO (targeting 4 sites in left striatum) and systemic administration of RLuc8.6 (150 mg/kg). The magnified view of the FUS-targeted region is shown on the left, and the contralateral site is shown on the right. The scale bar corresponds to 100 μm. **c)** Quantification of average fluorescence intensity of RLuc8.6 over the FUS-targeted region and the contralateral site. Data are presented as mean ± s.e.m. *n* = 4 mice for each group; \*\**P* < 0.01, using ratio paired t test (two-tailed). **d)** *In vivo* bioluminescent imaging representative data at field of view A (3.9 cm across). FUS targeting one site at left striatum was followed by an IV injection of RLuc8.6 (8 mg/kg) or PBS buffer control. RLuc8.6 alone group was intravenously injected with the same dose (8 mg/kg) of RLuc8.6 without FUS-BBBO procedure. After 1-hour, bioluminescent imaging was performed after administration of CTZ (i.p., 3.5 mg/kg). **e)** Quantification of *bioluminescence average radiance* (p/s/cm^2^/sr) at field of view C (13 cm across) in the head region 1h after FUS-BBBO. Data are presented as mean ± s.e.m. *n =* 6 mice for RLuc8.6 alone and FUS + RLuc8.6 groups, *n =* 3 for FUS alone group; \*\**P* < 0.01, \*\*\**P* < 0.001, ns (not significant), using one-way ANOVA with Tukey’s HSD test.

Collectively, these data revealed that the enzyme retains activity in the brain following RAID protocol, but the activity decays over the next 24 hours.

### Protein engineering improves the retention of delivered enzyme

We hypothesized that the enzyme’s clearance out of the brain interstitium led to the reduction of signal over time and decided to engineer RAID enzymes that can bind to the cells in the brain to improve the enzymes’ retention (**Fig. 3a**). To achieve this, we fused RLuc8.6 to engineered cell-adhesive peptides that mimic the extracellular matrix (ECM). These peptides have been shown to promote attachment to specific cells targeting cell-surface receptors^62-64^. To improve the enzyme retention after delivery and brain-tissue specificity, we fused the RLuc8.6 to IKVAV, GRGDS, and YIGSR, that are known to bind to neurons^65-67^. To avoid the skull attenuating the bioluminescent signal^68,69^, we analyzed the retention of each construct post-mortem in 2mm thick unfixed tissue sections. We tested two time points, 2 and 7 days after FUS-BBBO procedure (**Fig. 3b**) and found that FUS-mediated delivery led to over 10-fold higher level of bioluminescence over no-FUS control group (**Fig. 3c-e** and **Supplementary Fig. 3**), with all tested enzyme variants showing significant improvement in retention at 2 days (**Fig. 3d**, *n* = 4/5 mice per group, *P* = 0.0006, 0.0251 and 0.0387 for IKVAV, YIGSR and GRGDS fused RLuc8.6 respectively, One-way ANOVA with Tukey’s HSD test). Additionally, we found that fusing RLuc8.6 with one of the ECM-mimicking peptides (IKVAV) enhanced enzyme activity 2 days after FUS-BBBO delivery when compared to unmodified RLuc8.6 at 2-days (**Fig. 3d**). Fusion of Rluc8.6 and IKVAV^65^ (RLuc8.6-IKVAV) showed highest mean activity improvement at 2.3(±0.3)-fold over unmodified RLuc8.6 (*n* = 5 mice per group, *P* = 0.0452, One-way ANOVA with Tukey’s HSD test). The RLuc8.6-IKVAV was also the only RAID enzyme that was significantly different than RLuc8.6 control at 7-days after FUS-BBBO (**Fig. 3e**). We concluded that IKVAV fusion enhanced the RLuc8.6 retention, resulting in 113(±33)-fold higher level of bioluminescence than the control group without applying FUS (*n* = 5 mice per group, *P* = 0.0012, Oneway ANOVA with Tukey’s HSD test). Additionally, the RLuc8.6 remained functional within the brain parenchyma even at the 7-day timepoint.

**Fig. 3:**
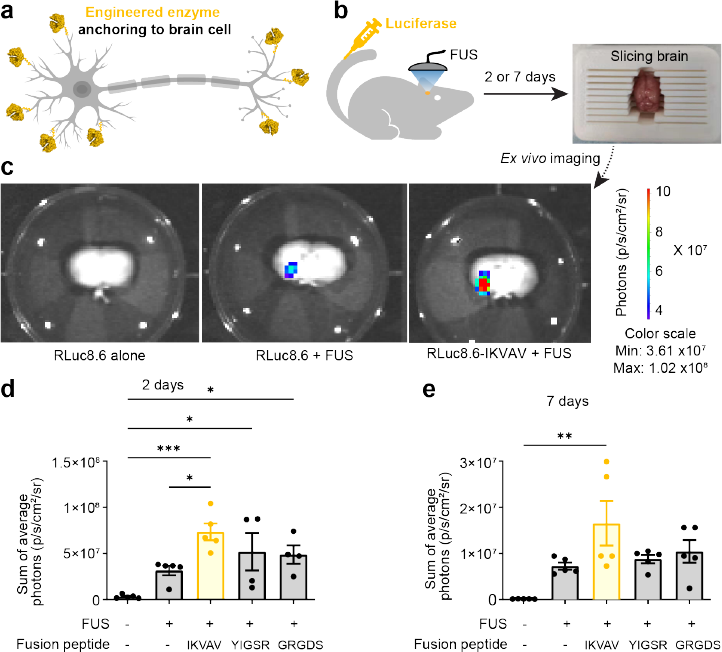
Engineering enzyme to improve retention in the brain. **a)** Schematic diagram of engineered cell-adhesive peptides that mimic the ECM for attaching prodrug enzymes to brain cells. **b)** Experimental scheme for measuring the retention of unmodified or engineered RLuc8.6 in the mouse brain. We conducted FUS-BBBO targeting three sites of left brain in mice following IV injection of RLuc8.6 (100 mg/kg) or engineered variant (104 mg/kg) and microbubbles. RLuc8.6 alone group was only intravenously injected with the same dose of RLuc8.6 without insonation. After 2 or 7 days, mouse brains were extracted without perfusion and cut into 2mm slices before bioluminescent imaging. **c)** *Ex vivo* bioluminescent imaging representative data for mice at 2 days after FUS-BBBO. After cutting, each 2mm brain section was individually transferred into a 6-well glass bottom plate filled with 2mL PBS buffer. BLI at field of view C (13 cm) was conducted immediately after adding 1mL dissolved CTZ with a final concentration of 10 μM. **d-e)** Sum of *bioluminescence average radiance* (p/s/cm^2^/sr) in all five sections of each mouse 2 days **(d)** or 7 days **(e)** after FUS-BBBO. Data are presented as mean ± s.e.m. *n =* 5 mice in each group except for RLuc-8.6-YIGSR and RLuc8.6-GRGDS groups at 2 days, *n =* 4. \**P* < 0.05, \*\**P* < 0.01, \*\*\**P* < 0.001, using One-way ANOVA with Tukey’s HSD test.

### Spatiotemporal modulation of dopamine receptor-expressing brain cells

After demonstrating that enzyme activity can be retained within the brain for multiple days, we next explored the feasibility of using RAID for localized neuromodulation. The concept of RAID can be theoretically applied to any enzyme-pro-drug pair as long as it uses a BBB-permeable prodrug. Thus, it is a flexible paradigm that is not dependent on targeting any specific molecular pathway. For our proof of concept, we chose a well-validated system with a clinically used prodrug and an enzyme that occurs naturally within the brain, and where any peripheral effects can be suppressed with a non-BBB permeable drug. Specifically, we chose the L-DOPA as a prodrug, which can be converted to the neurotransmitter dopamine through a protein enzyme, Aromatic-L-amino-acid decarboxylase (AADC)^70^. Dopamine is involved in multiple aspects of the brain function including motor control, reward, and motivation^71,72^. Since L-DOPA exists naturally in the healthy brain, AADC is expected to have some background activity, which theoretically could even allow for the enzyme delivery alone to have measurable effects. This system also holds high translational significance as it is relevant to the investigation of natural dopaminergic circuitry and has potential for the treatment of Parkinson’s Disease, where low amount of cellular AADC in basal ganglia limits dopamine production^73,74^. The peripheral effects of L-DOPA can be suppressed by administration of non-BBB-permeable carbidopa^75^, which is important to avoid peripheral effects. Thus, while AADC is naturally present in some brain regions, we hypothesized that locally increasing AADC concentration could sensitize the targeted brain region to lower doses of L-DOPA. Such sensitization could lead to a distinguishable site-specific neuromodulation that can be used in, for example, investigating the dopaminergic circuitry, testing the suitability of the brain region for AADC-expressing gene therapy, or lowering the dose of L-DOPA necessary to treat PD and thus the associated side effects. This approach would provide a proof-of-concept for the RAID paradigm, and a tool to evaluate effects of local perturbations of the dopaminergic network. We chose to activate dopamine receptor-expressing brain cells in striatum, which is a therapeutically relevant brain region already targeted for gene therapy for AADC deficiency^76^ and Parkinson’s disease^77^. To improve AADC’s retention within the brain, we fused it with IKVAV (**Fig. 3d-e**). Purified AADC-IKVAV (250 mg per kg of body weight) was injected intravenously. Then FUS-BBBO was targeted to a single site in the left caudate putamen (CPu) of wild-type mice. After 48 h, mice received an intraperitoneal (IP) injection of L-DOPA (2 mg/kg) and two hours later were perfused to evaluate spatial precision and presence of AADC-IKVAV delivery in tissue sections (**Fig. 4a**).

**Fig. 4:**
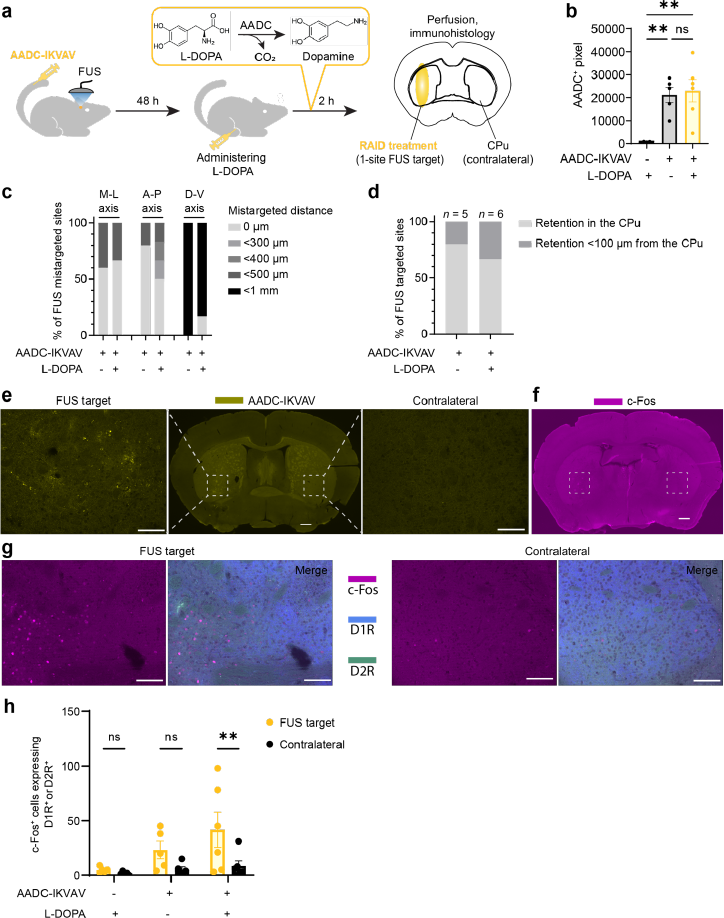
Localized activation of dopaminergic cells with engineered AADC. **a)** Experimental scheme for assessing control of cellular signaling with RAID. AADC-IKVAV (250 mg/kg) or PBS buffer was injected IV and was followed with FUS-BBBO targeting left Caudate Putamen (CPu). After 2 days, we administered L-DOPA (2 mg/kg, i.p.) or vehicle, and 2h later perfused the mice to analyze AADC-IKVAV retention and neuronal activation as indicated by c-Fos expression in tissue sections. **b)** Quantification of the AADC^+^ pixels in the FUS-targeted region among different groups. (Data are presented as mean ± s.e.m. *n =* 5 mice for FUS alone + L-DOPA and FUS + AADC-IKVAV groups, *n =* 6 mice for FUS + AADC-IKVAV + L-DOPA; \*\**P* < 0.01, ns (not significant), using One-way ANOVA with Tukey’s HSD test. One mouse in the FUS + AADC-IKVAV group was excluded due to no observed AADC-IKVAV retention. **c)** Spatial targeting precision of AADC-IKVAV delivery based on its histological retention. The representative immunostaining images of coronal sections from each mouse treated with FUS-mediated AADC-IKVAV delivery are shown in **Supplementary Fig. 4**. The estimation of distance from the intended target was made by comparison to the target programmed within the atlas embedded in the RK-50 ultrasound device. Displacement of less than 300 μm distance was considered accurate. *n =* 5 mice for FUS + AADC-IKVAV and *n =* 6 mice for FUS + AADC-IKVAV + L-DOPA group. M-L represents the medio-lateral axis, A-P represents the anterior–posterior axis, and D-V represents the dorso-ventral axis. **d)** Quantitative summary of AADC-IKVAV retention in the CPu within the FUS-targeted region. If there was no retention in the CPu, the distance of retention to the nearest CPu region was estimated using the same methodology as in point c). **e)** Representative immunostaining images of a coronal section from a mouse treated with FUS + AADC-IKVAV + L-DOPA for AADC staining. The rectangular areas within the FUS target and contralateral site were imaged at a higher magnification to compare AADC^+^ pixel counts. The scale bars correspond to 500 μm (for the entire coronal section) and 100 μm (for the magnified view), respectively. **f-g)** Representative immunostaining images of a coronal section adjacent to the area with the highest AADC-IKVAV retention in a mouse treated with FUS + AADC-IKVAV + L-DOPA, showing c-Fos (purple), dopamine receptor D1 (blue), and D2 (green) staining. Rectangular areas within the FUS target and the contralateral site were magnified for the comparison of c-Fos^+^ cells with or without the expression of dopamine receptors D1^+^ or D2^+^. The scale bars correspond to 500 μm **(f)** and 100 μm **(g)**, respectively. **h)** Quantification of c-Fos^+^ cells expressing dopamine receptors D1^+^ or D2^+^ in the FUS-targeted region, compared with contralateral site. Data are presented as mean ± s.e.m. *n =* 5 mice for FUS alone + L-DOPA and FUS + AADC-IKVAV group, *n =* 6 mice for FUS + AADC-IKVAV + L-DOPA; \*\**P* < 0.01, ns (not significant), using Two-way ANOVA with Sidak’s multiple comparison test.

We observed that AADC-IKVAV was retained at the FUS-targeted site following FUS-BBBO delivery, and administration of L-DOPA had no effect on that retention (**Fig. 4b-e and Supplementary Fig. 4)**. Specifically, mice treated with AADC-IKVAV, with or without subsequent L-DOPA administration, exhibited locally elevated AADC levels that were 38.5 (±6.1)-fold and 41.8 (±8.9)-fold higher, respectively, compared to the FUS alone control group (**Fig. 4b**, *n =* 5 mice for FUS + L-DOPA group and FUS + AADC-IKVAV group, and *n =* 6 mice for FUS + AADC-IKVAV + L-DOPA group; *P* = 0.0056 and *P* = 0.0021, respectively, One-way ANOVA with Tukey’s HSD test). Additionally, we assessed the spatial targeting precision of AADC-IKVAV delivery based on the site of AADC delivery as compared to the targeted site in a brain atlas embedded within the RK-50 ultrasound device. Among the 11 mice receiving AADC-IKVAV, 64% achieved the FUS target alignment (**Fig. 4a**) in both medio-lateral and anterior–posterior axes (**Fig. 4c**) with displacement of less than 300 microns as measured between the center of a simulated FUS beam profile in the histological section image and a brain atlas. For mice showing mistargeting, the displacement ranged from 300 to 1000 μm in any axis. This resulted in detectable AADC-IKVAV retention within the CPu for 73% of tested mice, while the remaining 27% showed efficient retention within 100 μm of the CPu (**Fig. 4d**).

To evaluate whether RAID with AADC and L-DOPA can elicit local cellular signaling, we first focused on the target cells that express dopamine receptors D1 and D2 that are found in the CPu^78^. We found a significant increase in c-Fos in cells showing D1- or D2-positive immunostaining in mice that were subjected to the full RAID treatment (**Fig. 4f-h and Supplementary Fig. 5**). Specifically, we saw 6.6 (±1.7)-fold higher count in the FUS-targeted region compared to the contralateral site (**Fig. 4h**, *n* = 6 mice, *P* = 0.0052, as determined by a Two-way ANOVA with Sidak’s multiple comparison test). No significant change was observed in the control groups, either those treated with FUS alone + L-DOPA (*n* = 5 mice, *P* = 0.9845, based on a Two-way ANOVA with Sidak’s multiple comparison test) or those treated with FUS + AADC-IKVAV without L-DOPA (*n* = 5 mice, *P* = 0.2046, as determined by a Two-way ANOVA with Sidak’s multiple comparison test). These results proved that RAID can specifically modulate activity of dopamine-responsive cells, and the effects were specific to the FUS-targeted hemisphere.

**Fig. 5:**
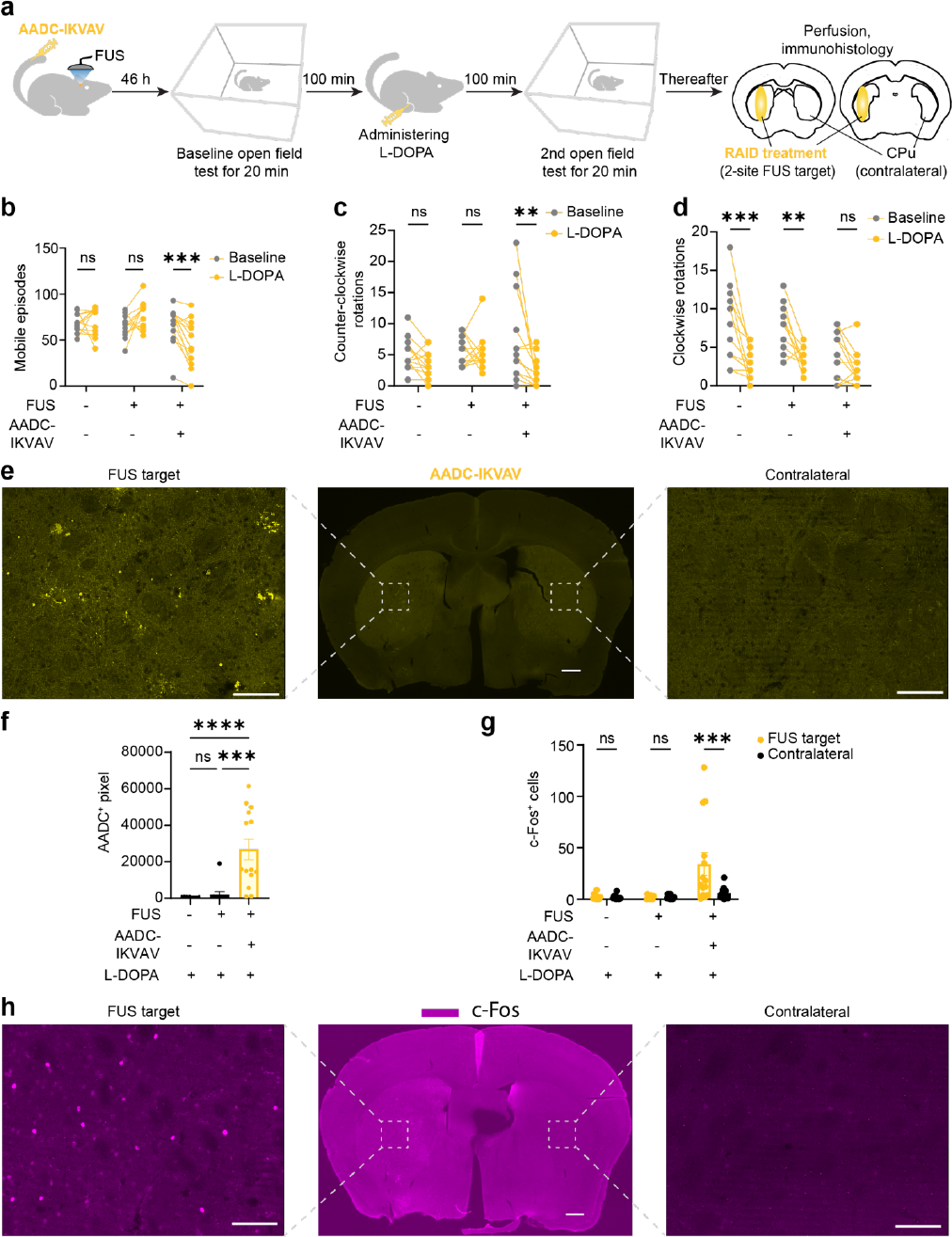
RAID control of behavior through targeted neuromodulation. **a)** Experimental scheme for RAID-induced control of motor behavior by targeted activation of dopamine neurons. We used FUS in two sites of the left striatum to deliver AADC-IKVAV (250 mg/kg, i.v. injection) or PBS buffer control. After 2 days, open field test (OFT) was performed twice before and after a single injection of the prodrug L-DOPA (2 mg/kg, i.p.), allowing 200 minutes between each test. The locomotor activity was recorded for 15 min after a habituation period of 5 min in each test. After the testing, we perfused mice 2h after L-DOPA administration for histologic examination of AADC-IKVAV retention and neuron activation (c-Fos^+^) to confirm localized neuromodulation. **b)** Mobile episodes, **c)** counterclockwise and **d)** clockwise rotations were measured in the open field test before and after administration of L-DOPA. Data are presented as mean ± s.e.m. *n =* 12 mice for wild-type group, *n =* 11 mice for FUS alone group and *n =* 14 mice for FUS + AADC-IKVAV group; \*\**P* < 0.01, \*\*\**P* < 0.001, ns (not significant), using Two-way ANOVA with Sidak’s multiple comparison test. **e)** Representative immunostaining images of a coronal section from a mouse treated with FUS + AADC-IKVAV + L-DOPA for AADC staining. Rectangular regions within both the FUS target and contralateral site were captured at a higher magnification to compare AADC^+^ pixel counts. The scale bars correspond to 500 μm (for the full coronal section) and 100 μm (for the enlarged view). **f)** Quantification of the AADC^+^ pixels in the FUS-targeted region (located at the left CPu) among different groups. Data are presented as mean ± s.e.m. *n =* 12 mice for wild-type + L-DOPA group, *n =* 11 mice for FUS + L-DOPA group and *n =* 14 mice for FUS + AADC-IKVAV + L-DOPA group; \*\*\**P* < 0.001, \*\*\*\**P* < 0.0001, ns (not significant), using One-way ANOVA with Tukey’s HSD test. **g)** Quantification of c-Fos^+^ cells in the FUS targeted region at the CPu, compared with contralateral site. Data are presented as mean ± s.e.m. *n =* 12 mice for wild-type + L-DOPA group, *n =* 11 mice for FUS + L-DOPA group and *n =* 14 mice for FUS + AADC-IKVAV + L-DOPA group; \*\*\**P* < 0.001, ns (not significant), using Two-way ANOVA with Sidak’s multiple comparison test. **h)** Representative immunostaining images of a coronal section adjacent to the area with the highest AADC-IKVAV retention in a mouse treated with FUS + AADC-IKVAV + L-DOPA, showing c-Fos staining in purple. Rectangular areas within the FUS target at the CPu and the contralateral site were magnified to compare c-Fos^+^ cell counts. The scale bars correspond to 500 μm (for the entire coronal section) and 100 μm (for the magnified view), respectively

While we focused FUS-BBBO on CPu, some amount of AADC delivery was also present in off-target areas, as would be expected given the size of FUS-BBBO in our equipment (ovoid with ∼1 × 5mm major axes diameters). These regions are highly interconnected with neighboring and midbrain neurons^78^, allowing for potential secondary activation or inhibition of neuronal activity. However, we found no significant changes in the group of mice that received only FUS-BBBO without AADC-IKVAV delivery (**Supplementary Figure 6a**., *n* = 5 mice, *P* = 0.9989, based on a Two-way ANOVA with Sidak’s multiple comparison test). We also found no statistically significant changes in these regions in mice receiving FUS + AADC-IKVAV alone (*n* = 5 mice, *P* = 0.0832, based on a Two-way with Sidak’s multiple comparison ANOVA test), or FUS + AADC-IKVAV + L-DOPA (*n* = 6 mice, *P* = 0.0571, based on a Two-way ANOVA with Sidak’s multiple comparison test).

**Fig. 6:**
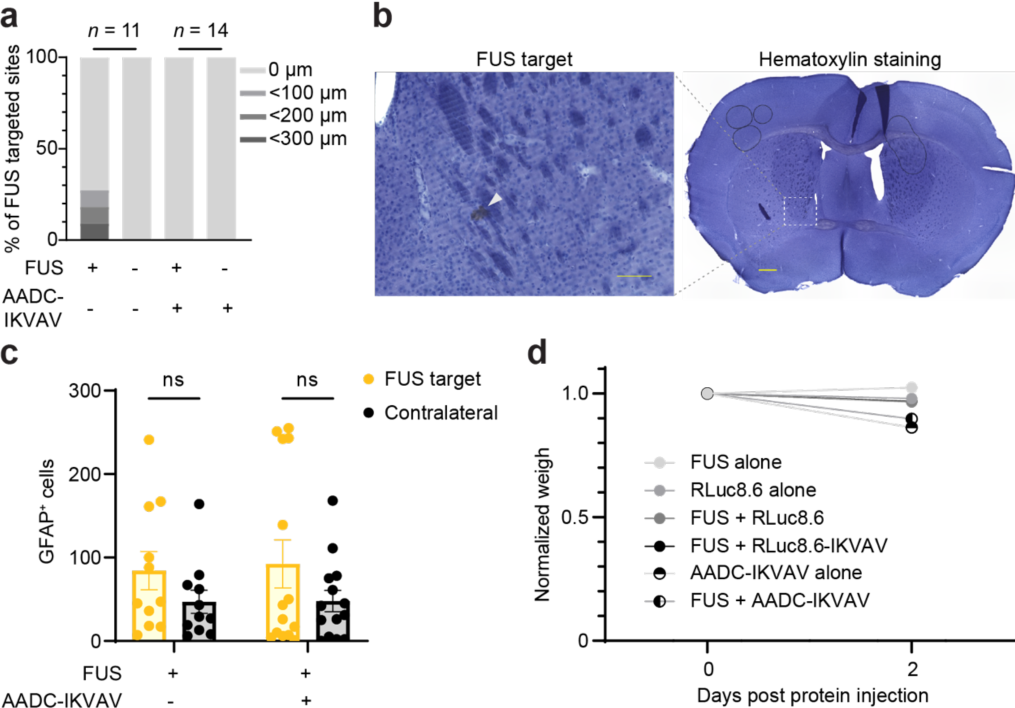
Safety assessment of RAID protocol in behavior analysis. **a)** Histological classification of brain tissue damage at 25 FUS targeted sites in 25 mice following FUS-BBBO with or without AADC-IKVAV. The adjacent sections of those used for c-Fos staining (**Fig. 5h**) were used for hematoxylin staining. *n =* 11 mice for FUS alone group and *n =* 14 mice for FUS + AADC-IKVAV group. **b)** Representative hematoxylin-stained brain section containing a small area of hemorrhage (indicated by the white arrow) in a mouse treated with FUS alone. Scale bars correspond to 100 μm (left) and 500 μm (right), respectively. **c)** Quantification of GFAP^+^ cells in the FUS targeted region at striatum, compared with contralateral site. Data are presented as mean ± s.e.m. *n =* 11 mice for FUS alone group and *n =* 14 mice for FUS + AADC-IKVAV group; ns (not significant), using Two-way ANOVA test. **d)** Mouse body weight analysis. The weight just before FUS-BBBO or intravenous injection of unmodified or engineered RLuc8.6 and AADC-IKVAV was used to normalize the weight of each mouse during the experiment. Data are presented as mean ± s.d. *n =* 11 mice for FUS alone group, *n* = 10 mice per unmodified or engineered RLuc8.6 group, *n =* 6 mice for AADC-IKVAV alone group and *n =* 14 mice for FUS + AADC-IKVAV group. Intra-group differences between day 0 and 2 days after protein injection were identified using the Two-way ANOVA with Sidak’s multiple comparison test. The statistical results were summarized in **Supplementary Table 1**.

### Behavioral validation of RAID

To measure the effects of RAID neuromodulation on behavior, we focused on targeting dorsal striatum with two FUS sites, to increase the amount of on-target delivery of AADC in the CPu region that contains dopamine-responsive cells^79,80^. Dopamine is involved in regulation of motor behavior^81,82^, and previous studies indicated that localized, unilateral, delivery of dopamine to striatum resulted in changes to locomotor activity^83-85^ and rotational behavior^86,87^. To evaluate whether RAID could induce similar behavioral changes after the prodrug administration, we used our proof-of-concept pair of AADC-IKVAV and L-DOPA.

We delivered AADC-IKVAV via FUS-BBBO to 2 sites in the left striatum region of wild type mice. After 46 h mice were placed in an open field (OF) to test their locomotor activity for 20 minutes (**Fig. 5a**). Mice were then placed in the home cages, and 100 minutes later received L-DOPA (2 mg/kg) via IP injection and were tested again in OF.

We found that mice with RAID treatment showed 38±8% decrease in mobile episodes, where mice move after an extended period of freezing (longer than 2 s) (**Fig. 5b**, *n* = 14 mice, *P* = 0.0005, Two-way ANOVA with Sidak’s multiple comparison test), in line with previous studies showing that direct unilateral injection of dopamine into the striatum led to a reduction in locomotor activity (e.g., moving, walking)^83,88^. However, no significant change in episodes of mobility was observed in the control groups treated with either FUS alone or wild type mice. Second, we assessed whether RAID could induce contralateral rotations, as previously demonstrated with direct unilateral dopamine injections into the caudate putamen^87,89^. We found that L-DOPA indeed reduced the counter-clockwise (ipsilateral) rotations by 63±10% in mice with full RAID treatment (**Fig. 5c and Supplementary movies 1-2**, *n* = 14 mice, *P* = 0.0018, Two-way ANOVA with Sidak’s multiple comparison test) but not in the control mice with FUS alone or no treatment at all (*n =* 11 mice for FUS alone group, *n =* 12 mice for wild-type group; *P* = 0.9992 and 0.6019 respectively, Two-way ANOVA with Sidak’s multiple comparison test). In contrast, the clockwise (contralateral) rotation showed an opposite trend. The untreated and FUS alone groups showed decreased clockwise rotation after injection of L-DOPA by 52±8% and 44±13% (**Fig. 5d**, *n =* 12 mice for wild-type group, *n =* 11 mice for FUS alone group; *P* = 0.0001 and *P* = 0.0011 respectively, Two-way ANOVA with Sidak’s multiple comparison test), but RAID treatment prevented that decrease (*n* = 14 mice, *P* = 0.2951, Two-way ANOVA with Sidak’s multiple comparison test). To evaluate whether the AADC-IKVAV alone had any effects on mouse behavior, we compared the motor behavior of mice in the open field before L-DOPA administration. We found that AADC-IKVAV had no effect on the number of mobile episodes, or counter-clockwise rotations, but interestingly we did find significant effect on clockwise rotations (**Supplementary Fig. 7**, *P* = 0.0128 and *P* = 0.0181 for WT group vs full RAID treatment, and FUS-alone vs full RAID treatment, respectively; One-way ANOVA with Tukey’s HSD test). There were no significant effects in any test for the FUS-alone treatment compared to the wildtype mice. Overall, these results suggested that RAID could control behavior following the L-DOPA administration. The results were consistent with the expected mechanism of action of dopamine delivered unilaterally to the CPu.

Immediately after the behavioral testing, we perfused the mice and extracted brains for histological analysis. Immunostaining against the delivered enzyme showed that AADC-IKVAV was present in the FUS-targeted area in all mice treated with RAID (**Fig. 5e and Supplementary Fig. 8**), with an average of 93.3(±19.5)-fold and 13.3(±2.8)-fold higher levels of local AADC compared to the untreated and FUS alone groups, respectively (**Fig. 5f**, *n =* 12 mice for wild-type group, *n =* 11 mice for FUS alone group and *n =* 14 mice for FUS + AADC-IKVAV + L-DOPA group; *P* < 0.0001 and *P* = 0.0002 respectively, One-way ANOVA with Tukey’s HSD test). Additionally, RAID-treated mice exhibited a 5.7(±1.8)-fold increase in ipsilateral c-Fos-positive cells (**Fig. 5g-h**, *n* = 14 mice, *P* = 0.0004, Two-way ANOVA with Sidak’s multiple comparison test), whereas no significant changes were observed in untreated mice or mice treated with FUS alone (*n =* 12 mice for wild-type group, *n =* 11 mice for FUS alone group; *P* = 0.9998 and *P* > 0.9999 respectively, Two-way ANOVA with Sidak’s multiple comparison test). Taken together, our data suggests that RAID was effective in control of specific behaviors and was able to induce site-specific brain activity. Thus, RAID could be a useful non-surgical method for investigating behavioral effects of localized drug action.

### Safety evaluation

To assess the safety of the RAID approach, we stained brain sections of the mice in behavior study with hematoxylin and anti-GFAP (glial fibrillary acidic protein) antibody respectively. Hematoxylin staining showed that the RAID approach didn’t cause observable tissue damage or bleeding (**Fig. 6a**). Out of 25 mice treated with FUS only 3 (12%) showed any detectable tissue damage. In those mice, the damage was scattered (**Fig 6b**) and any damage present was contained within the area of radius between 100 and 300 μm (**Fig 6b**), in line with previous dicated that there were no significant changes in GFAP^+^ astrocytes observed in the ipsilateral region following FUS-BBBO procedure alone (**Fig. 6c**, *n* = 11 mice, *P* = 0.2328, Two-way ANOVA with Sidak’s multiple comparison test;). The RAID treatment, similarly, did not affect the numbers of GFAP+ astrocytes in the targeted area when compared to the contralateral site (**Fig. 6c and Supplementary Fig. 9**, *n* = 14 mice, *P* = 0.0828, Two-way ANOVA with Sidak’s multiple comparison test).

Throughout the duration of the RAID protocol, we conducted an analysis of the body weight of mice. The group treated with FUS + RLuc8.6 and FUS + RLuc8.6-IKVAV experienced a weight loss of 3.4±0.7% and 3.2±0.8% respectively after 2 days following protein injection (*n* = 10 mice for each group, *P* = 0.0071 and *P* = 0.0118 respectively, Twoway ANOVA with Sidak’s multiple comparison test). Additionally, this minor weight loss was temporary, and mice began to regain body weight three days after protein injection, with no significant weight loss was observed on day 3 compared to day 0 (**Supplementary Fig. 10 and Supplementary Table 2**, *n* = 5 mice for both FUS + RLuc8.6 and FUS + RLuc8.6-IKVAV groups, with P-values of 0.9521 and 0.6775, respectively, according to the Two-way ANOVA with Sidak’s multiple comparison test). A systemic injection of AADC-IKVAV exhibited a significant and larger decrease in body weight of 10.3±0.5% within 2 days of FUS-BBBO (**Fig. 6d and Supplementary Table 1**, *n* = 14 mice, *P* < 0.0001, Two-way ANOVA with Sidak’s multiple comparison test). In contrast, mice subjected to FUS alone did not display any notable changes in body weight (*n* = 11 mice, *P* = 0.0974, Two-way ANOVA with Sidak’s multiple comparison test). In a separate control group of mice, we administered AADC-IKVAV intravenously without utilizing FUS to evaluate peripheral effects of the AADC enzyme, resulting in a considerable reduction in body weight of 13.8±0.8% within 2 days post-injection (*n* = 6 mice, *P* < 0.0001, Two-way ANOVA with Sidak’s multiple comparison test). Overall, the RAID approach showed no tissue damage or bleeding, and no significant changes in astrocytes, indicating safety to the brain tissue. The weight loss, while present, was associated with systemic administration of AADC and its peripheral effects that can be suppressed with non-BBB-permeable carbidopa^75^.

## DISCUSSION

In this study, we demonstrated a proof-of-concept of a noninvasive, non-genetic, site-specific neuromodulation, which we call Regionally Activated Interstitial Drugs, or **RAID**. Our approach uses FUS-BBBO to deliver an engineered enzyme that attaches to the brain interstitium and equips the targeted site with ability to produce drugs from BBB-permeable, systemically-supplied inert prodrugs. RAID has unique advantages that can be used to study brain activity or in therapy planning. First, RAID is non-genetic, thus reducing the concerns of immunogenicity of viral vectors and trouble with their readministration^20,21^. Second, RAID uses delivery of proteins with FUS-BBBO, which can be delivered using safe ultrasound pressures^92^, with protein enzymes being well below the limit of the particle size that can be delivered with FUS-BBBO^93^. Third, RAID allows for neuromodulation in the absence of BBB opening, unlike in the case FUS-BBBO-based delivery of small molecule drugs^33,36,37^ where drug presence is inherently linked with the opened BBB presenting a confounding factor to neuromodulation. RAID allows for a single FUS-BBBO procedure to provide multi-day long neuromodulation, as opposed to several hours while the BBB stays open^38,39^. Lastly, unlike in the case of long-term drug release from nanoparticles, RAID is designed to tune the magnitude of neuromodulation in the brain by simply changing the dose of the prodrug. Similarly, by increasing the dose of the prodrug, one can compensate for the loss of enzyme over time, providing stable degree of drug activation over time.

In our proof of concept, RAID was sufficient to induce changes in localized neuronal activity, and to modulate behavior with a specific pair of an enzyme and prodrug. However, RAID is a versatile concept. It is compatible with any enzyme that can turn a prodrug into an active drug, be delivered to the brain with FUS-BBBO and effect on-demand localized drug action. Our results demonstrated the basic capabilities of RAID in mice with localized delivery and multi-day retention of activity for two example enzymes.

A single injection of L-DOPA significantly activated cells expressing either of the tested dopamine receptors (D1 and D2) in the vicinity of delivered AADC-IKVAV. We have also observed expected behavioral effects, that were consistent with previously published work^81-87^. Unexpectedly, we found that even wild-type mice showed a degree of lateralized rotations. This could be caused by the asymmetrical environment of our custom-made open field arenas (e.g., uneven light reflection, fan noise), in accordance with previous studies showing that environmental context influences open-field behavior^94-96^, especially when combined with the habituation leading to lower overall movement of mice when placed in the open field the second time^97,98^. Importantly, our negative controls (FUS alone group) behaved indistinguishably from wild-type mice and the changes in rotation behavior induced by RAID treatments were consistent with previous studies involving unilateral intracranial dopamine injections into the GPu^87,89^, and that our negative controls (FUS alone group) behaved indistinguishably from wild-type mice.

While we did not find significant effects of the AADC delivery alone on c-Fos accumulation or on two of the three covariates tested in the open field, we did find an effect of AADC on clockwise rotations (**Supplementary Fig. 7c**). One possibility is that in wild-type mice AADC can convert endogenous L-DOPA into dopamine and introduce background activation of neurons in the caudate putamen^80,99^. This effect would likely be reduced in Parkinson’s disease, where endogenous L-DOPA levels are lower^100,101^ and where AADC/L-DOPA RAID would be most useful as a therapeutic strategy. While we only observed significant effect in one of the tested scenarios, it is possible that multiple factors, such as asymmetry of the rotations in the open field, residual background levels of L-DOPA, or other effects coincided in that scenario to produce the isolated significant effect. On the other hand, if the administration of an enzyme alone can indeed produce lasting behavioral effects dependent on the background conversion of endogenous neurotransmitters, it would present a useful tool for interrogation of neural circuitry or therapy. Specifically, for those neurotransmitters that do not have a BBB-permeable prodrug, administration of RAID enzyme alone can provide localized neuromodulation, albeit one that will not be tunable on-demand with varied prodrug levels. In the intended application of RAID as a tool for investigation of circuitry or therapy planning, the most critical aspect of RAID is that induces localized neuromodulation that results in a distinguishable behavioral readout after the treatment, with or without the prodrug administration.

With that in mind, we envision RAID could be useful in a number of scenarios including, for example, large animal studies, as a noninvasive and reversible alternative to lesioning or deep-brain stimulation. Without the need of surgical resection or viral vector admnistration, multiple brain regions could be studied on different days using RAID. Without genetic component, RAID would not result in induction of neutralizing antibodies against the viral vectors^20-22^. In therapy planning, for example, RAID could be used to validate the presumed seizure focus, before resection^102^, genetic modification for chemogenetic neuromodulation e.g., with Acoustically Targeted Chemogenetics (ATAC)^27^, or invasive deep brain stimulation^103^, with use of a combination of drugs^104^ and enzymes^105,106^ that silence neuronal activity. An unacceptable side effect profile of such silencing, or lack of effects on seizure reduction over several-day long RAID neuromodulation could help refine the site or cell-type targeted for treatment.

Further improvements to RAID could include testing other cell-adhesive peptides to promote attachment of delivered enzymes to the surrounding brain cells^107,108^, to improve the enzyme’s retention in the brain. In our study, we observed that enzyme retention was enhanced through fusing with ECM-mimicking peptide that binds to neurons. This led to a 1-week long retention of RLuc8.6-IKVAV in the brain after a single session of FUS-BBBO delivery. Another approach could be to tether RAID enzymes to ECM, which has previously been used to capture and retain the secreted ECM proteins^109^. Other improvements could include improving the tissue specificity of the delivered enzymes, to ensure lower exposure of peripheral tissues to the enzyme. In its current, proof-of-concept, form RAID allows for studying effects of localized neuromodulation. However, just as any other systemically administered drug therapy, RAID is prone to peripheral effects that will need to be considered in each application. Lastly, a third area of potential improvement would be development of more sensitive enzymes that can respond to lower concentrations of prodrugs and thus reduce any possible prodrug side effects. As an additional advantage, more sensitive enzymes would allow for reduction of the enzyme dose, thus reducing the cost and potential immune response.

We chose to evaluate the proof-of-concept of RAID paradigm with the use of AADC and L-DOPA to control dopamine receptor-expressing brain cells. This approach simultaneously has advantages and disadvantages. On one hand, use of a naturally occurring enzyme in the body reduces concerns of immune response against the enzyme, and use of a clinically approved drug, L-DOPA, enables a path of translation. This approach allows for prodrug-induced site-specific effects of neuromodulation, forming a good tool to plan therapy or study the effects of site-specific neuromodulation. On the other hand, AADC itself may have side effects due to its ability to synthesize other neurotransmitters (such as serotonin) and trace amine neuromodulators including phenylethylamine, tyramine, and tryptamine^110,111^. AADC-IKVAV might catalyze the above decarboxylation reactions utilizing the endogenous aromatic L-amino acid substrates after delivery through the BBB. Additionally, these biological reactions could also occur peripherally after the systemic administration of AADC-IKVAV, potentially leading to weight loss due to the significant roles played by peripheral dopamine and serotonin in metabolic regulation^112,113^. These effects will need to be taken into consideration and managed during any potential treatment or experiment involving this specific enzyme and prodrug pair.

To mitigate potential side effects associated with naturally occurring enzymes and neurotransmitters, the develoment of a more specific and orthogonal enzyme-prodrug pair would enable more precise neuromodulation. Another option is a reduction of systemic exposure of RAID enzymes. For example, focused ultrasound-mediated intranasal brain drug delivery (FUSIN)^114^ could be used for site-specific enzyme delivery, and be followed-up with systemic administration of a prodrug to lower exposure to an enzyme in major organs. Implementing strategies like FUSIN can enhance the targeted delivery of prodrug enzymes and minimize potential systemic side effects. In the current form, RAID can be useful in therapy planning and site-specific neuromodulation to investigate brain circuitry. Further improvement of each component of RAID will enable control of different cell-types and molecular pathways with greater duration, lowered side effects, and improved specificity.

## MATERIALS AND METHODS

### Animals

Wild-type C57BL/6J mice (12-18 weeks of age) were purchased from Jackson Laboratory and housed in a 12 h light/dark cycle and were provided with water and food ad libitum. All experiments were performed under a protocol approved by the Institutional Animal Care and Use Committee of Rice University.

### Plasmid construction

For constructing the recombinant protein expression plasmid of RLuc8.6, the protein coding sequence (CDS) of RLuc8.6 was amplified from pcDNA-RLuc8.6-535^59^ (Addgene ID 87125) and subcloned into the vector pRSETb^115^ (Addgene ID 89536) with a N-terminal His-tag through Gibson assembly at the BamHI and EcoRI site. The CDSs of ECM-mimicking peptides (synthesized by GenScript) were assembled to the C-terminal of RLuc8.6 for creating the expression plasmids of engineered variants. Similarly, the expression plasmid of AADC-IKVAV was constructed by subcloning AADC amplified from Sino Biological plasmid (HG29995-CF) and IKVAV fragment into the above vector using the same restriction sites. Primers used for cloning are listed in **Supplementary Table 3**. CDSs, noncommercial plasmid sequences, and subcloning insertion sites are listed in **Supplementary Table 4**.

### Protein expression and purification

The recombinant protein was expressed in *Escherichia coli* and purified by Ni-affinity chromatography. For RLuc8.6 and engineered variants, *Escherichia coli* BL21 (DE3) cell was transformed with expression construct respectively and grown in Terrific Broth (TB) medium at 37°C to an OD_600_ of ∼0.6 before induction with 0.1 mM IPTG at 20°C for overnight. Harvested cell pellets from 1-liter cultures were resuspended in 40 mL ice-cold lysis buffer (50 mM sodium phosphate, 300 mM NaCl, 10 mM imidazole, 10% glycerol, pH 8.0) for sonication. The supernatant after centrifugation at 17, 500 RPM, 4°C for 45 min was loaded into the glass chromatography columns (Bio-Rad, catalog number 7372522) and incubated with Ni-NTA agarose resin (Qiagen, catalog number 30210) on ice for 1h. The column was washed and eluted with a stepwise imidazole gradient (10 mM to 500 mM) of lysis buffer through gravity flow. Eluates were concentrated with a Corning® Spin-X® UF 20 mL centrifugal filter unit (10 kDa cutoff), and then buffer exchanged into PBS using the PD-10 desalting column. Eluted proteins were concentrated again, analyzed by SDS-PAGE, and quantified by a NanoDrop spectrophotometer. Similarly, AADC-IKVAV was expressed in *Escherichia coli* BL21 (DE3) cell and purified by Ni-affinity chromatography through gravity flow as described above, except that the over-expression was induced by 0.05 mM IPTG at 22 °C and it was gradually buffer changed into PBS while being concentrated with a 30 kDa cutoff filter.

### FUS-BBBO

C57BL/6J male mice (12-18 weeks of age) were anaesthetized with 2.5% isoflurane in 1.5% O_2_ and shaved on top of skull using a trimmer. A catheter, made by a 30-gauge needle connected to PE10 tubing, was inserted into the tail vein, affixed in place using tissue glue and then flushed with 10 units (U) mL^−1^ of heparin in sterile saline (0.9% NaCl). Subsequently, the mouse was mounted on the RK50 (FUS Instruments) stereotactic platform using the ear bars and bite bar/nose cone. A midline scalp incision was vertically made to expose the skull after disinfecting the site using three alternating scrubs of chlorhexidine scrub and chlorhexidine solution. The locations of Lambda and Bregma were registered in the RK50 software using the guild pointer. Next, sterile ultrasound gel was applied on the surface of skull before placing the ultrasound transducer in a tank, both of which were filled with degassed water. The mice were then sequentially injected via tail vein with purified recombinant protein in PBS buffer and approximately 1.5 × 10^6^ DEFINITY microbubbles (Lantheus) per gram of body weight diluted in sterile saline. Immediately after injections of protein and microbubbles, the mice were insonated using RK-50 FUS system with axial and lateral diameter of 5 mm and 1.2 mm, respectively. FUS target coordinates used for each experiment are listed in **Supplementary Table 5**. The ultrasound parameters used were 1.5 MHz, 10 ms duration, 1000 ms burst period for 120 pulses. The pressure at 0.3 MPa was used for FUS-BBBO based on preliminary tests in our lab, except that 0.36 MPa was chosen for the experiments in **Fig. 2** according to previous studies^116,117^. Following insonation, the mice were placed back to home cage for recovery after closing the scalp incision with tissue glue.

### Immunohistochemical analysis of RLuc8.6

Mice (*n* = 4) were injected with RLuc8.6 (20 mg/mL, 150 mg/kg, i.v.) right before FUS-BBBO targeting 4 sites at left striatum with 2 min interval between insonations. After 1h, the mice were sacrificed by transcardial perfusion with cold heparinized (10 U mL^−1^) PBS following induction of anaesthesia using ketamine/xylazine solution (80 mg/kg and 10 mg/kg, respectively), and immediately afterwards with 10% neutral buffered formalin. The brains were extracted and postfixed for 24-48 hours in the same fixative at 4°C before being sliced into 50 μm coronal sections using a vibratome (Leica). The slices were blocked in 10% normal goat serum (SouthernBiotech) and 0.3% Triton-X solution in PBS for 1h at room temperature and then incubated with a primary rabbit anti-RLuc antibody (1:1000, PA1-180, ThermoFisher) in blocking buffer for overnight at 4°C. Subsequently, the sections were washed three times (15min each) in PBS and then incubated with a secondary goat anti-rabbit antibody conjugated to Alexa Fluor 647 (1:500, A-21245, ThermoFisher) in blocking buffer for 2h at room temperature. After washing three times (15min each) in PBS, these sections were mounted onto glass slides using the mounting media (Vector Laboratories) with DAPI and allowed to air-dry overnight in dark prior to imaging.

### *In vivo* BLI

Two groups of mice (*n* = 6 mice per group) underwent FUS-BBBO targeting one site at left striatum immediately after i.v. injection of RLuc8.6 (2 mg/mL, 8 mg/kg) or equivalent volume of PBS buffer. The third group of mice (*n* = 3) was injected intravenously with the same dose of RLuc8.6 without FUS-BBBO procedure. BLI was conducted 1h, 24h, 48h and 96h after FUS insonation with an IVIS spectrum imager (Perkin Elmer). The mice were injected with CTZ (2.5 mg/mL, 3.5 mg/kg, i.p.; cat# 303-INJ, Nanolight Technology) after being anaesthetized with 2.5% isoflurane in 1.5% O_2_. Bioluminescence images under similar anesthesia were taken every 5min until luminescent signal of the head peaked, usually 5-15 min after injection of CTZ. The BLI parameters used were open filter for emission, automatic exposure time (mostly 5s), aperture (f/stop) 1, binning 8, field of view A (3.9 cm, imaging the head) and C (13 cm, imaging the whole body). The bioluminescence signal was quantified by calculating the average radiance (p/s/cm^2^/sr) in the head region of view C imaging using the Living Image software (Caliper Life Sciences).

### *Ex vivo* analysis of engineered RLuc8.6

FUS was performed to target three sites in the left hemisphere of C57BL6J mice (n = 5 mice per group) immediately after systemic administration of RLuc8.6 (20 mg/mL, 100 mg/kg) or engineered variant (20 mg/mL, 104 mg/kg) and microbubbles by tail vein injection. The dose of engineered RLuc8.6 was increased accordingly based on their molecular weight for injecting the same number of molecules as unmodified RLuc8.6. The control group (*n* = 5 mice) was injected intravenously with the same dose of unmodified RLuc8.6 without FUS. The mice were euthanized using CO_2_ without perfusion at two time points (2-day and 7-day) after FUS-BBBO procedure. Immediately afterwards, the brains were extracted, washed with ∼20 mL PBS buffer in a 50mL conical tube for 30s, and then cut into 2mm sections without olfactory bulb and brainstem using a coronal Slicer (Invitrogen). The sections were individually transferred into a 6-well glass bottom plate (Cellvis) filled with 2mL PBS buffer. BLI was performed using an IVIS spectrum imager (Perkin Elmer) immediately after adding 1mL dissolved CTZ (cat# 303-INJ, Nanolight Technology) with a final concentration of 10 μM. The parameters used here were similar as *in vivo* BLI with 1s of exposure time and field of view C. The average radiance (p/s/cm^2^/sr) of each brain section was quantified with the Living Image software (Caliper Life Sciences) and summed to compare the activity retention of RLuc8.6 among different groups. At timepoint of 2 days, two mice (each from RLuc-8.6-YIGSR and RLuc8.6-GRGDS group respectively) showed FUS-BBBO-related tissue damage, resulting abnormally high levels of bioluminescence, and thus were excluded from analysis.

### c-Fos activation with engineered AADC

Three groups of mice underwent FUS-BBBO procedure to target a single site at left striatum immediately after intravenous injection of recombinant protein AADC-IKVAV (20 mg/mL, 250 mg/kg) or equivalent volume of PBS buffer (*n* = 5 mice for FUS alone + L-DOPA group). After 48h, the mice were given a single dose of L-DOPA (0.1 mg/mL, 2 mg/kg, i.p.; Spectrum Chemical) 10min after injection of carbidopa (1 mg/mL, 25 mg/kg, i.p.; Sigma-Aldrich), both of which were dissolved in sterile saline containing 2.5 and 0.125 mg/mL ascorbic acid. The FUS + AADC-IKVAV control group (*n* = 6 mice) did not receive L-DOPA injection, while the experimental group (*n* = 6 mice for the FUS + AADC-IKVAV + L-DOPA group) received L-DOPA injection. After 2 hours, the mice were euthanized by transcardial perfusion, and their brains were extracted and sliced into 50 μm coronal sections following 24-48 hours of fixation.

Immunostaining of AADC was performed as follows: (1) incubate the brain sections in 1X antigen retrieval solution (catalog number: 00-4955-58; Invitrogen) overnight in a 60°C water bath; (2) block sections in 5% normal goat serum (SouthernBiotech) and 0.3% Triton-X solution in PBS for 1h at room temperature; (3) incubate with primary rabbit anti-AADC antibody (1:500, 10166-1-AP, Proteintech) for staining AADC-IKVAV in blocking buffer for 2h at room temperature; (4) after washing three times in PBS (15min each, the same as below), incubate with secondary goat anti-rabbit antibody conjugated to Alexa Fluor 647 (1:750, A-21245, ThermoFisher) in blocking buffer for 2h at room temperature; (5) after washing three times, mount sections onto glass slides using the mounting media (Vector Laboratories) with DAPI and air-dry overnight in dark before imaging.

The section displaying the strongest AADC fluorescence was selected as the representative for each mouse, and its adjacent section was stained separately to identify c-Fos and dopamine receptor D1 and D2 positive cells. The procedure was carried out as follows: (1) block sections in 10% normal goat serum (SouthernBiotech) and 0.3% Triton-X solution in PBS for 1h at room temperature; (2) incubate with primary rabbit anti-c-Fos (1:2000, 2250S, Cell Signaling Technology) antibody in PBS with 0.3% Triton-X for 2h at room temperature; (3) after washing three times in PBS (15min each, the same as below), incubate with primary anti-D1 Dopamine Receptor (1:500, D2944, MilliporeSigma) and anti-Dopamine Receptor D2 (1:500, Cat. # 376 205, Synaptic Systems) antibodies in 5% normal goat serum and 0.3% Triton-X solution in PBS overnight at 4 °C; (4) after washing three times, incubate with secondary goat anti-rabbit antibody conjugated to Alexa Fluor 647 (1:500, A-21245, ThermoFisher) in PBS with 0.3% Triton-X for 2h at room temperature; (5) after washing three times, incubate with secondary goat anti-rat antibody conjugated to Alexa Fluor 546 (1:500, A-11081, ThermoFisher) and goat anti-Guinea Pig antibody conjugated to Alexa Fluor 488 (1:500, A-11073, ThermoFisher) in 5% normal goat serum and 0.3% Triton-X solution in PBS for 2h at room temperature; (6) after washing three times, mount sections onto glass slides using the mounting media (Vector Laboratories) with DAPI and air-dry overnight in dark before imaging.

### Locomotor behavior test

Two groups of mice underwent FUS-BBBO targeting two sites at left striatum immediately after intravenous injection of recombinant protein AADC-IKVAV (*n* = 14 mice for FUS + AADC-IKVAV group, 20 mg/mL, 250 mg/kg) or equivalent volume of PBS buffer (*n* = 11 mice for FUS alone group). Another group without FUS-BBBO procedure served as wild type control (*n* = 12 mice for WT group). After 46h, the first session of behavioral test was performed in a custom-made non-transparent open field box (30.5 cm X 30.5 cm) as a baseline. Each mouse was individually placed into the apparatus center with a light intensity and background noise at ∼320 lux and 46 dB (400Hz peak), respectively. After a habituation period of 5 min, the locomotor activity of a freely-moving mouse was recorded for 15min by a video tracking system (Stoelting Co.) consisting of an overhead camera connected to a computer with Any-Maze software. Behavioral measures include total distance traveled, average and maximum speed, freezing time and episodes, mobile/immobile time and episodes, clockwise and counter-clockwise rotations and head turn angles. The mouse was returned to the home cage immediately after testing. After 90min, the mice were intraperitoneally injected with carbidopa (25 mg/kg) and 10min later L-DOPA (2 mg/kg) as described above. We conducted the second session of open field test 100min after L-DOPA injection.

Immediately afterwards, the mice were sacrificed for immunohistochemical analysis of AADC-IKVAV retention as already mentioned. The section displaying the strongest AADC fluorescence was selected as the representative for each mouse, and its adjacent section was stained separately to identify c-Fos positive cells. The procedure was carried out as follows: (1) block sections in 10% normal goat serum (SouthernBiotech) and 0.3% Triton-X solution in PBS for 1h at room temperature; (2) incubate with primary rabbit anti-c-Fos (1:2000, 2250S, Cell Signaling Technology) and chicken anti-tyrosine hydroxylase (1:2000, SKU: TYH, Aves Labs) antibodies in PBS with 0.3% Triton-X for 2h at room temperature; (3) after washing three times in PBS (15min each, the same as below), incubate with primary mouse Anti-6X His tag antibody (1:10000, ab18184, Abcam) in blocking buffer overnight at 4 °C; (4) after washing three times, incubate with secondary goat anti-rabbit antibody conjugated to Alexa Fluor 647 (1:500, A-21245, ThermoFisher) and goat anti-chicken antibody conjugated to Alexa Fluor 488 (1:500, A-11039, ThermoFisher) in PBS with 0.3% Triton-X for 2h at room temperature; (5) after washing three times, incubate with secondary goat anti-mouse antibody conjugated to Alexa Fluor 546 (1:1000, A-21143, ThermoFisher) in blocking buffer for 2h at room temperature; (6) after washing three times, mount sections onto glass slides using the mounting media (Vector Laboratories) with DAPI and air-dry overnight in dark before imaging.

### Safety analysis

To investigate potential lesions resulting from the use of FUS-BBBO in the RAID protocol, we performed hemotoxicity staining (*n* = 11 mice for FUS alone + L-DOPA group and *n* = 14 mice for FUS + AADC-IKVAV + L-DOPA group) on the adjacent sections of each representative section used for c-Fos staining, which was associated with behavior analysis as shown in **Fig. 5**. Hemotoxicity staining was conducted according to the following steps: (1) The section was immersed in 100% ethanol for 1 minute; (2) It was then soaked in 95% ethanol for 1 minute; (3) Subsequently, a 1-minute wash in H2O was performed; (4) The section was soaked in 50% hematoxylin (H&E Staining Kit, Abcam ab245880) for 1 minute; (5) This was followed by a 3-minute wash in H2O; (6) A 20-second soak in Bluing Reagent (H&E Staining Kit, Abcam ab245880) was carried out; (7) Another 3-minute wash in H2O followed; (8) Finally, the sections were mounted onto glass slides using mounting media (Vector Laboratories) and air-dried overnight in the dark before imaging.

We also stained those adjacent sections separately with an anti-GFAP antibody to examine astrocytic activation. The staining procedure involved the following steps: (1) The sections were blocked in a solution of 10% normal goat serum (SouthernBiotech) and 0.3% Triton-X in PBS for 1 hour at room temperature; (2) They were then incubated with primary mouse GFAP antibody conjugated to Alexa Fluor 647 (1:500, sc-33673 AF647, Santa Cruz Biotechnology) in a blocking buffer for 2 hours at room temperature; (3) Following three washes in PBS (15 minutes each), the sections were mounted onto glass slides using mounting media (Vector Laboratories) with DAPI and left to air-dry overnight in the dark before imaging.

To assess the potential impact of the RAID approach on the body weight of the mice involved in behavior analysis (*n* = 11 mice for the FUS alone + L-DOPA group and *n* = 14 mice for the FUS + AADC-IKVAV + L-DOPA group), we recorded their body weight on a daily basis before and after FUS-BBBO administration. Additionally, we included another group of mice (*n* = 6 mice) for comparison, which received intravenous injections of the same dose of AADC-IKVAV (20 mg/mL, 250 mg/kg) but did not undergo any FUS-BBBO treatment. Their body weight was also recorded in a similar manner; however, we unintentionally omitted weighing 4 of them on the day following the AADC-IKVAV injection. Similarly, we also recorded the mouse body weight before and after the intravenous injection of unmodified or engineered RLuc8.6, which is related to the study presented in **Fig. 3**.

### Histological imaging

All histological images were obtained using a fluorescence microscope (BZ-X810, Keyence). To evaluate RLuc8.6 retention, as depicted in **Fig. 2b** and **Supplementary Fig. 1**, we acquired fluorescence images of striatal sections using a ×4 objective in DAPI (nuclei, shown in blue) and far-red (RLuc8.6, shown in purple) channels. Subsequently, we identified a 3x3 binned area within the FUS target region of the representative section in the left striatum, exhibiting the highest RLuc8.6 retention. This specific area was then imaged using a ×40 objective, while also capturing images of the corresponding contralateral areas in the right striatum.

To analyze AADC-IKVAV retention, as depicted in **Fig. 4 and Fig. 5**, stained coronal sections within the striatum were captured using a ×4 objective in DAPI (nuclei, not shown) and far-red (AADC, shown in yellow) channels. Among these sections, the one displaying the most intense AADC fluorescence was chosen as the representative for each mouse. Subsequently, the selected sections were further examined at a higher magnification (×40 objective) for quantitative comparison. Specifically, we identified a rectangular area (5x9 binning each) within the FUS target region in the left striatum of each representative section, which exhibited the highest AADC-IKVAV retention. This area was imaged using a ×40 objective. Additionally, corresponding contralateral areas in the right striatum were also imaged.

To quantify the c-Fos-positive cells as shown in **Fig. 4**, we initially imaged the stained coronal sections using a ×4 objective in DAPI (nuclei, not shown), Cy3 (D1R, shown in blue), green (D2R, shown in green), and far-red (c-Fos, shown in purple) channels. Within the left striatum region, we selected specific 5x9 binned square areas corresponding to the highest observed AADC fluorescence in the adjacent AADC-stained section. Subsequently, we captured images of these areas using a ×40 objective. We also imaged the corresponding contralateral areas in the right striatum.

To quantify the c-Fos-positive cells related to **Fig. 5**, we initially imaged the stained coronal sections using a ×4 objective in DAPI (nuclei, not shown) and far-red (c-Fos, shown in purple) channels. Within the left striatum region, we selected specific 3x3 binned square areas corresponding to the highest observed AADC fluorescence in the adjacent AADC-stained section. Subsequently, we captured images of these areas using a ×40 objective. We also imaged the corresponding contralateral areas in the right striatum.

To assess the safety of the RAID protocol, as shown in **Fig. 6**, we initially imaged the hemotoxicity-stained brain sections using only the brightfield channel with a ×4 objective. Subsequently, sections exhibiting hemorrhage were further observed at a higher magnification (×40 objective). Additionally, the brain sections stained for GFAP were imaged using a ×4 objective in DAPI (nuclei) and far-red (GFAP, purple) channels. Following this, we selected three square areas in both the left and right striatum respectively, each with 3x3 binning, for imaging with a ×40 objective to quantify GFAP^+^ cells.

### Quantitative analysis of histology images

To quantify the c-Fos positive cells, as shown in **Fig. 4 and 5**, we analyzed the histological images using ZEISS ZEN software (Version 3.5). For both the DAPI (nuclei) and far-red (c-Fos, purple), we applied a Brightness (White) threshold of 6045 and a Gamma value of 1. The images were then overlayed, and the c-Fos^+^ cells were manually counted. Similarly, we manually counted the GFAP^+^ cells, as shown in **Fig. 6**, by applying a Brightness (White) threshold of 5686 and a Gamma value of 1 for both the DAPI (nuclei) and far-red (GFAP, purple) channels.

To quantify the AADC-IKVAV retention in the FUS targeted region, as shown in **Fig. 4b and Fig. 5f**, we analyzed the histological images using ImageJ (version 1.53t). Specifically, we calculated the average pixel intensity of the AADC-IKVAV channel image captured on the contralateral side of each representative section to establish the background for that section. Subsequently, we determined the number of positive pixels of the image captured on the FUS target side of that section, defined as pixels exhibiting an intensity value more than 3-fold higher than the background observed in the corresponding contralateral side image.

To manually count the c-Fos-positive cells in **Fig. 4h-i**, three scorers (Z.H., S.N., and R.J.) independently assessed all c-Fos staining sites. R.J. served as the blinded scorer, and the averaged counts from all three scorers were rounded for the final statistical analysis.

To address potential bias and validate our counting approach, all c-Fos staining sites from mice in **Fig. 5g** were independently counted by two scorers (A.M. and C.H.). C.H. acted as the blinded scorer. The comparisons among different groups yielded consistent trends. For instance, RAID-treated mice showed a *5*.*7(±1*.*8)-fold* increase (counted by A.M.) and a 22.3(±12.5)-fold increase (counted by C.H.) in ipsilateral c-Fos-positive cells compared to the contralateral site in the behavior study (*n* = 14 mice, *P* = 0.0006 and *P* = 0.0024, respectively, Two-way ANOVA test with Sidak’s multiple comparison test). The interexperimenter variability (average percentage difference) for the FUS + AADC-IKVAV + L-DOPA group was 9.3±2.1% for the FUS target (*n* = 14 mice, *P* = 0.7464, heteroscedastic two-tailed t-test) and 23.5±5.9% for the contralateral site (*n* = 14 mice, *P* = 0.268, heteroscedastic two-tailed t-test).

### Statistical analysis

The statistical analysis was performed using GraphPad Prism software version 9.0. All quantitative data were presented as mean ± s.e.m., except for normalized weight in **Fig. 6d**, mean ± s.d. A two-tailed ratio paired t-test was used for **Fig. 2c**. One-way ANOVA followed by Tukey’s honestly significant difference test was used for comparing the means of three or more independent groups. Two-way ANOVA followed by Sidak’s multiple comparison test was used to compare the mean differences between groups that are affected by two factors. The difference between groups was considered statistically significant when *p < 0.05, **p < 0.01, ***p< 0.001, ****p < 0.0001.

## ACKNOWLEDGEMENTS

The authors thank Dr. Gregory Corder (University of Pennsylvania) and Dr. Caleb Kemere (Rice University) for helpful discussions on the behavior results. This research was supported by the Welch Foundation grant to J.O.S #C-2048-20200401. Related work in the laboratory was also supported by the The G. Harold and Leila Y. Mathers Foundation, and the Michael J. Fox foundation for Parkinson’s Research.

## AUTHOR CONTRIBUTIONS

J.O.S. and Z.H. conceived and planned the research. Z.H. and J.O.S. designed the experiments and wrote the manuscript with input from all other authors. Z.H. performed and participated in all experiments described in this study. A.M. performed protein purification, *in vivo* and histological experiments, and cell counting. S.N. participated in the *in vivo* BLI and hematoxylin staining. J.P.S. and C.H. helped with histological experiments, and M.H. and R.J. assisted with FUS-BBBO and cell counting, respectively.

