## Supplementary Information for "Engineering Regionally-Activated Drugs for Neuroscience"

**Supplementary Figures 1-10, Tables 1-5 and Movies 1-2**

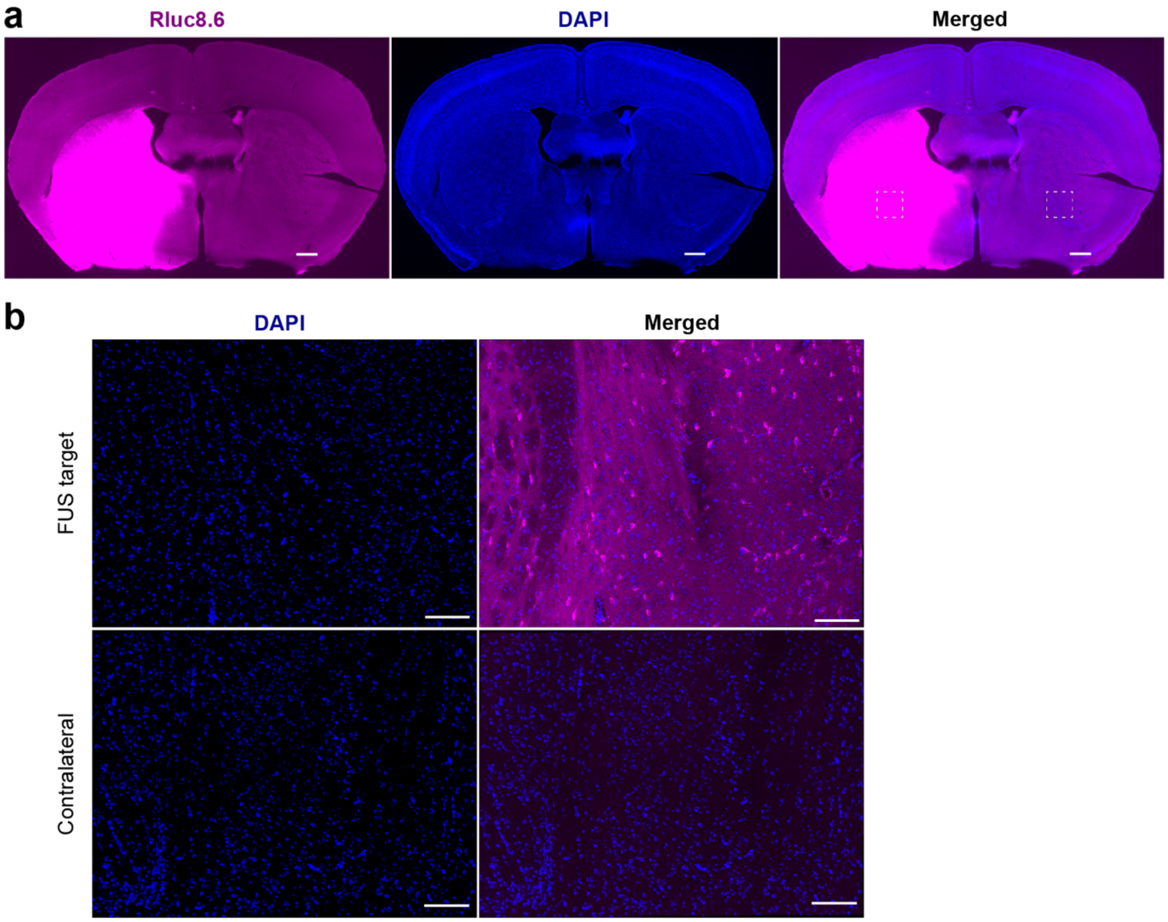

**Supplementary Figure 1. Additional representative immunostaining images after RLuc8.6 delivery related to Fig. 2b. a**) Fluorescence imaging of coronal section with RLuc8.6 retention (purple) in a mouse 1h after FUS-BBBO (targeting 4 sites in the left striatum) and systemic administration of RLuc8.6 (150 mg/kg), as shown in **Fig. 2b**. Cell nuclei were stained with DAPI (blue). The square areas of the FUS target and contralateral site in those sections were imaged at a greater magnification to compare the average fluorescence intensity, as shown in **Fig. 2c**. **b)** Magnified view of FUS targeted region (upper) and contralateral site (lower) corresponding to the square areas in **a**. Scale bars correspond to 500 μm (**a**) and 100 μm (**b**), respectively.

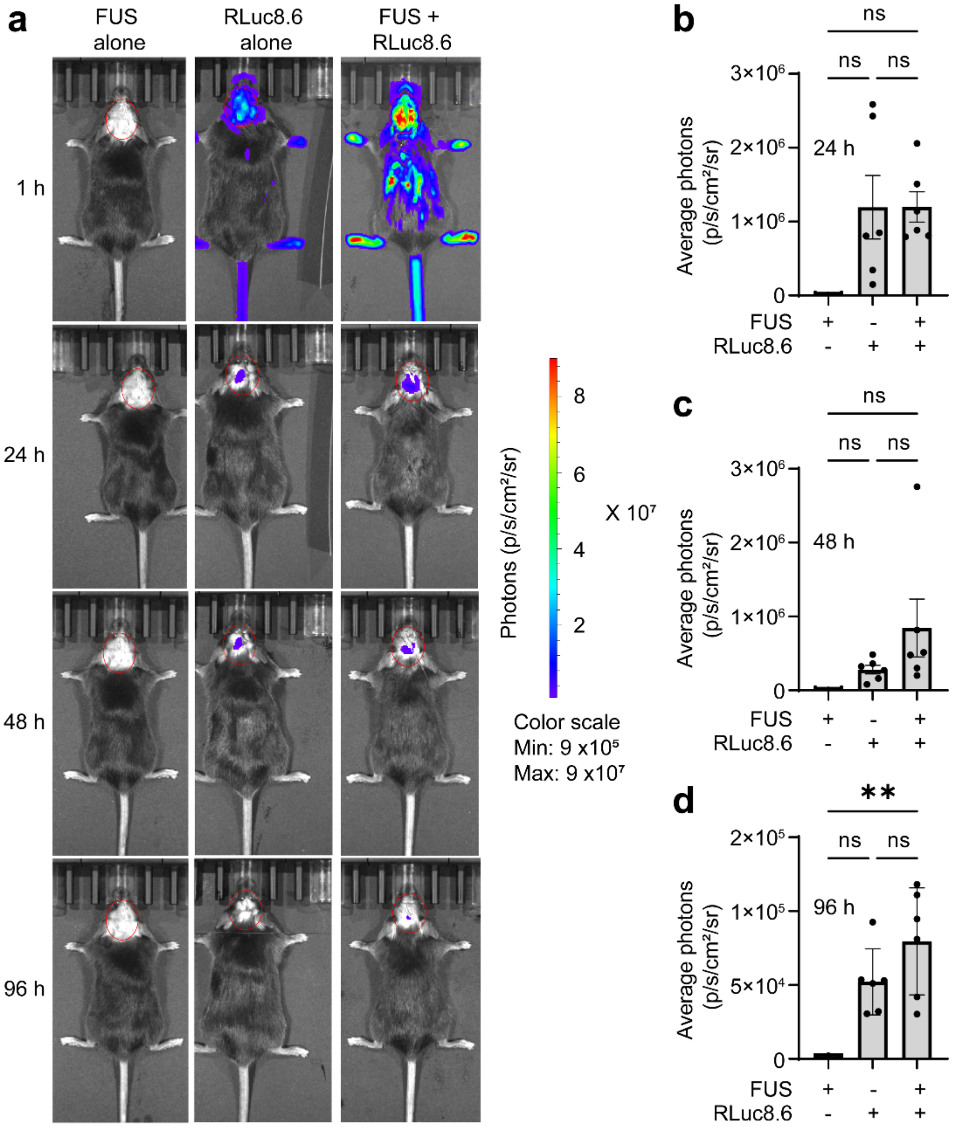

**Supplementary Figure 2. *In vivo* BLI at different time points after delivering RLuc8.6. a)** *In vivo* bioluminescent imaging representative data at field of view C (13 cm) for the mice shown in **Fig. 2d**, 1 h, 24 h, 48 h and 96 h after IV injection of RLuc8.6 (8 mg/kg) or PBS buffer with or without insonation. The mice were administered CTZ (i.p., 3.5 mg/kg) before each bioluminescent imaging. **b-d)** Quantification of bioluminescence average radiance (p/s/cm$^2$/sr) at field of view C (13 cm) in the head region 24 h, 48 h and 96 h after FUS-BBBO. Data are presented as mean ± s.e.m. $n$ = 6 mice for RLuc8.6 alone and FUS + RLuc8.6 groups, $n$ = 3 for FUS alone group; **$P$ = 0.0045, ns (not significant), using One-way ANOVA test.

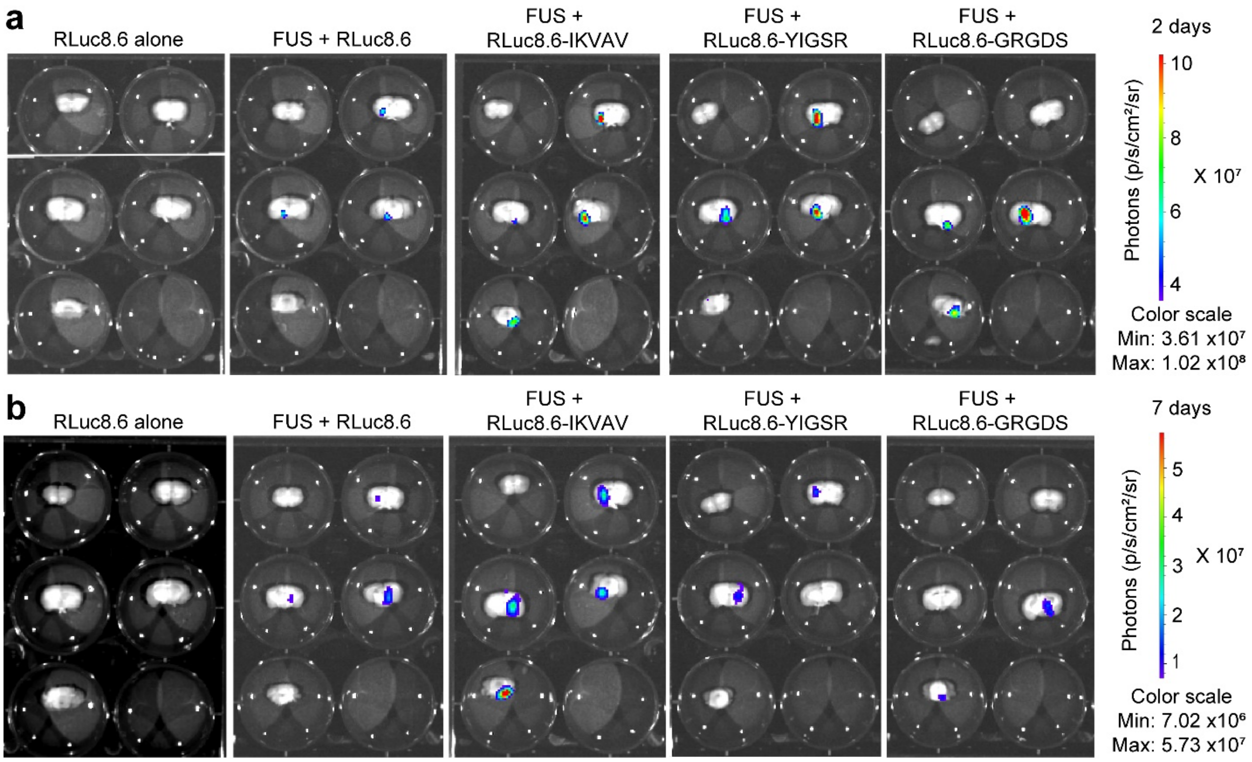

**Supplementary Figure 3. *Ex vivo* bioluminescent imaging (BLI) of brain sections for measuring unmodified or engineered RLuc8.6 retention after delivery. a-b** Additional *ex vivo* bioluminescent imaging representative data for mice at 2 days (**a**) or 7 days (**b**) after FUS-BBBO. We performed BLI at field of view C (13 cm across) immediately after adding 1mL dissolved CTZ with a final concentration of 10 μM.

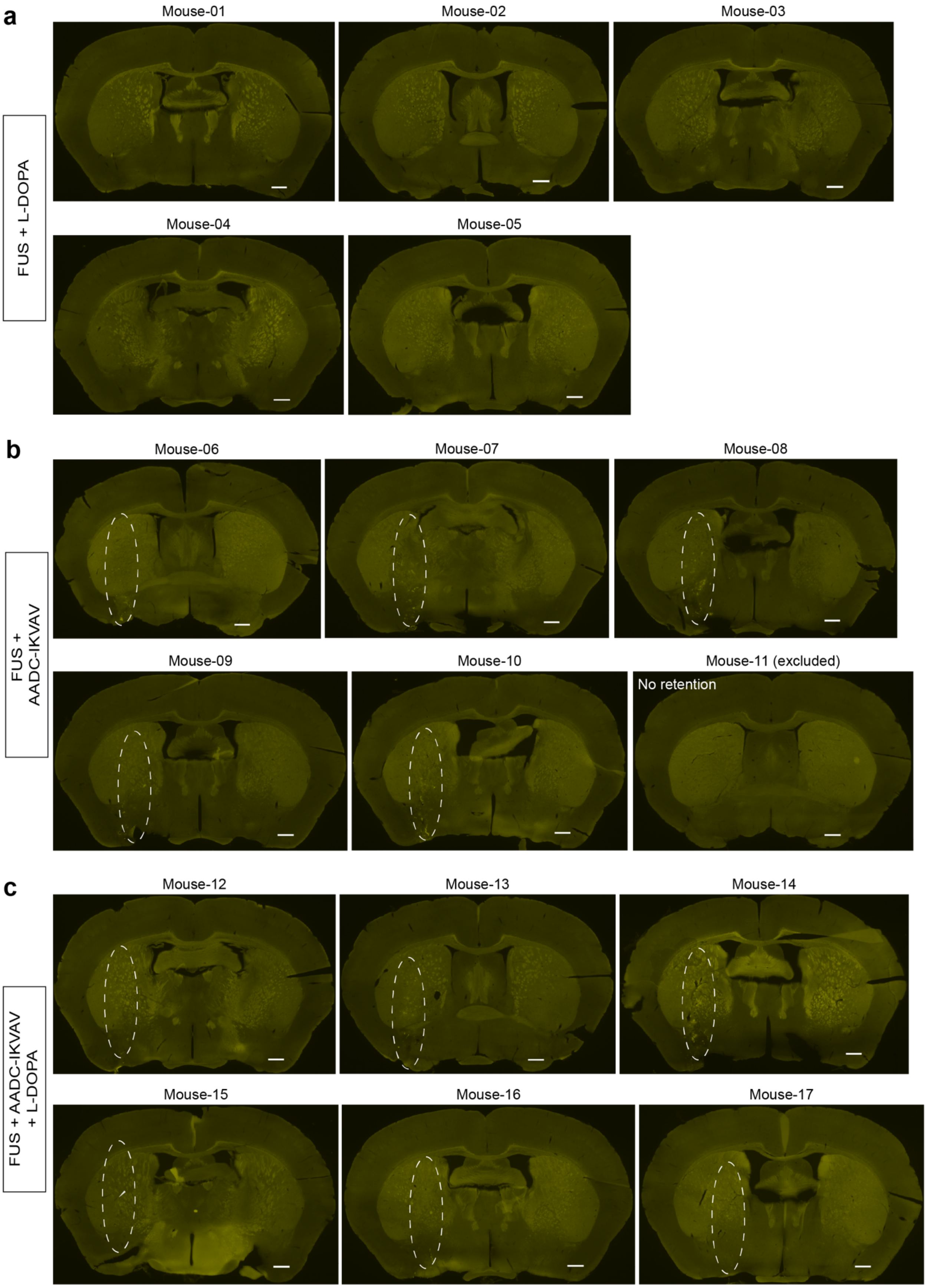

a
Mouse-01
Mouse-02
Mouse-03
FUS + L-DOPA
Mouse-04
Mouse-05
b
Mouse-06
Mouse-07
Mouse-08
FUS + AADC-IKVAV
Mouse-09
Mouse-10
Mouse-11 (excluded)
No retention
c
Mouse-12
Mouse-13
Mouse-14
FUS + AADC-IKVAV + L-DOPA
Mouse-15
Mouse-16
Mouse-17

**Supplementary Figure 4. Representative immunostaining images of AADC staining for all mice in the study related to Fig. 4.** Representative immunostaining images of coronal sections from mice treated with FUS + L-DOPA **(a)**, FUS + AADC-IKVAV **(b)**, or FUS + AADC-IKVAV + L-DOPA **(c)**, depicting AADC staining in yellow, are shown. These sections highlight the highest AADC staining signals around the striatum area in each mouse. Mouse-11 in the FUS + AADC-IKVAV group was excluded due to lower $AADC^+$ pixel counts compared to the FUS + L-DOPA group's average and no AADC-IKVAV retention. The scale bar corresponds to 500 μm. The presumed actual FUS target region for the mice with AADC-IKVAV retention was labeled using a theoretical ultrasound beam (represented by a white dashed line), and its size was estimated from the image scale bar and the dimensions of the ultrasound transducer, which had axial and lateral diameters of 5 mm and 1.2 mm, respectively. Mouse-15's AADC-IKVAV retention was indicated by a white arrow for easy identification.

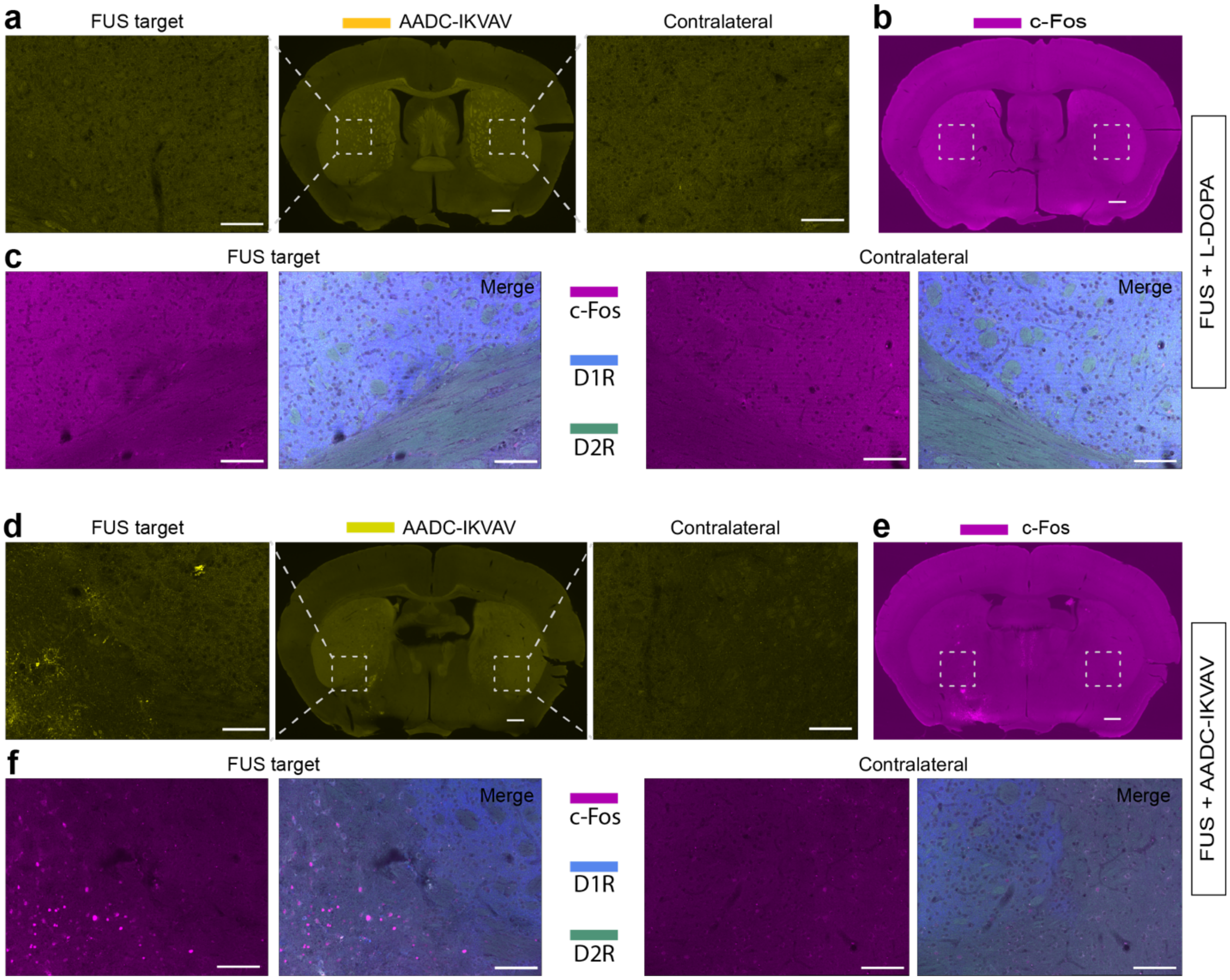

**Supplementary Figure 5. Representative immunostaining images of control groups related to Fig. 4. a and d) AADC staining:** Representative immunostaining images of a coronal section from a mouse treated with FUS + L-DOPA (**a**) or FUS + AADC-IKVAV (**d**) for AADC staining. The rectangular areas within the FUS target and contralateral site were imaged at a higher magnification to compare AADC$^+$ pixel counts. The scale bars correspond to 500 µm (for the entire coronal section) and 100 µm (for the magnified view), respectively. **b-c and e-f) c-Fos and dopamine receptor staining:** Representative immunostaining images of a coronal section adjacent to the area with the highest AADC-IKVAV retention in a mouse treated with FUS + L-DOPA (**b-c**) or FUS + AADC-IKVAV (**e-f**), showing c-Fos (purple), dopamine receptor D1 (blue), and D2 (green) staining. Rectangular areas within the FUS target and the contralateral site were magnified for the comparison of c-Fos$^+$ cells with or without the expression of dopamine receptors D1$^+$ or D2$^+$. The scale bars correspond to 500 µm **(b and e)** and 100 µm **(c and f)**, respectively.

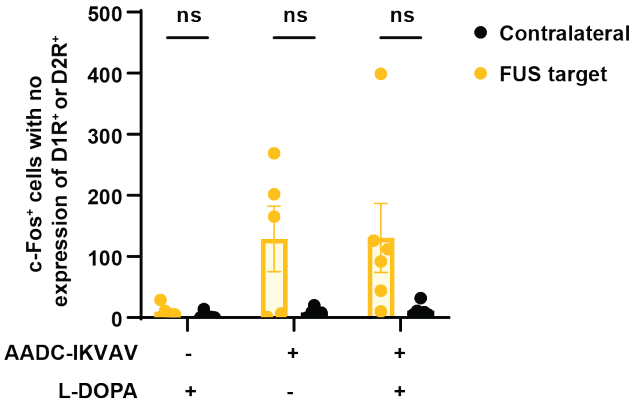

**Supplementary Figure 6**. Quantification of c-Fos$^+$ cells with no expression of dopamine receptors D1$^+$ or D2$^+$ in the FUS-targeted region, compared with contralateral site. Data are presented as mean ± s.e.m. *n* = 5 mice for FUS alone + L-DOPA and FUS + AADC-IKVAV group, *n* = 6 mice for FUS + AADC-IKVAV + L-DOPA; ns (not significant), using Two-way ANOVA test.

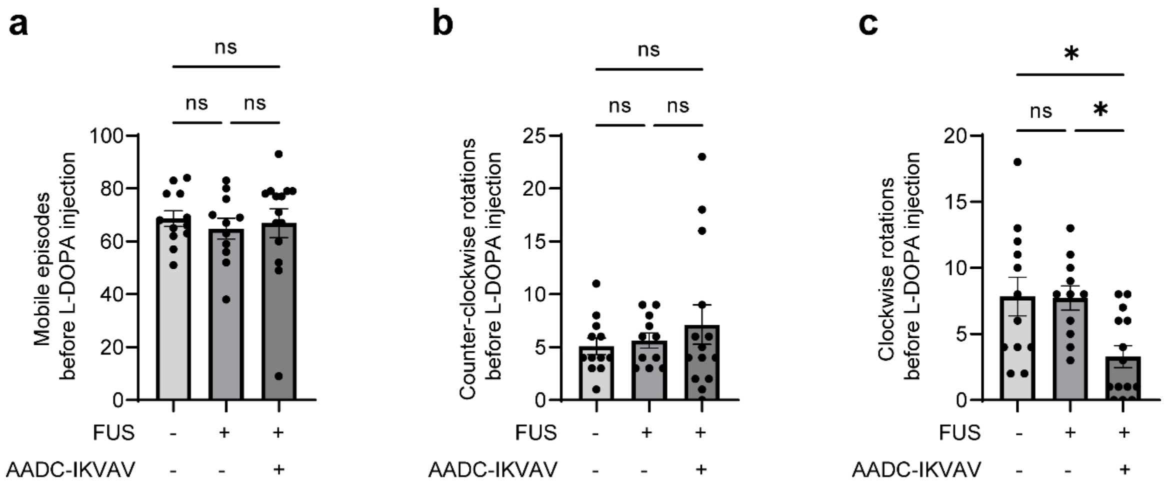

**Supplementary Figure 7. Mobile episodes (a), counter-clockwise rotations (b), and clockwise rotations (c) before the administration of L-DOPA were compared among groups.** Data are presented as mean ± s.e.m. *n* = 12 mice for wild-type group, *n* = 11 mice for FUS alone group and *n* = 14 mice for FUS + AADC-IKVAV group; **P* < 0.05, ns (not significant), using One-way ANOVA test.

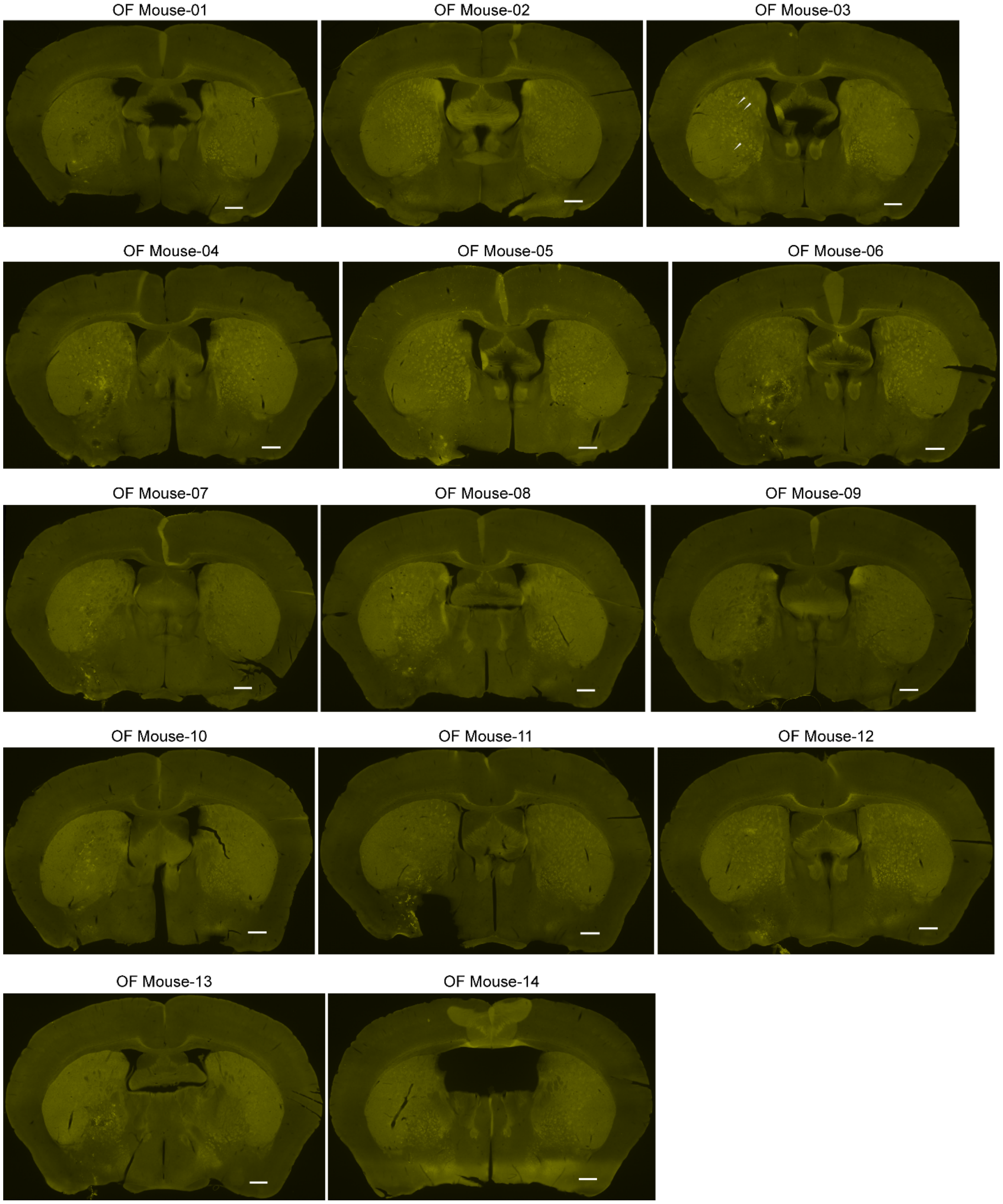

**Supplementary Figure 8. Representative immunostaining images of AADC staining for all mice with AADC-IKVAV delivery related to Fig. 5.** Representative immunostaining images of coronal sections from mice treated with FUS + AADC-IKVAV + L-DOPA after behavior tests, depicting AADC staining in yellow, are displayed. These sections highlight the highest AADC staining signals around the striatum area in each mouse. The scale bar corresponds to 500 μm. White arrows were used to indicate AADC-IKVAV retention in OF Mouse-03 for easy identification.

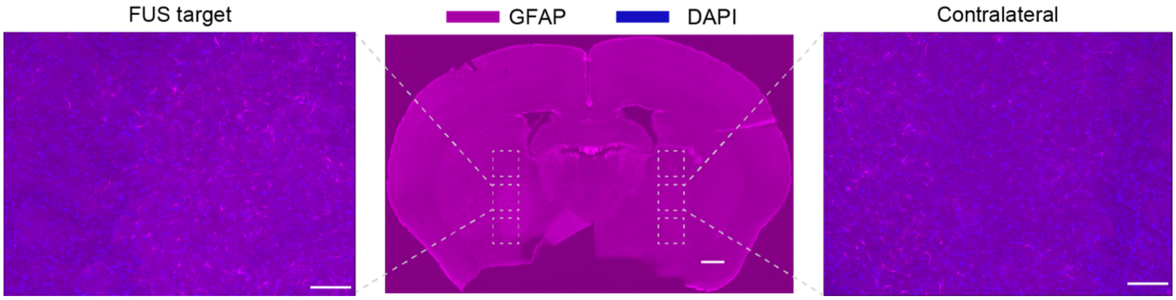

**Supplementary Figure 9. Representative immunostaining images for GFAP (purple) in a mouse treated with FUS and AADC-IKVAV.** Cell nuclei were stained with DAPI (blue). Middle image: fluorescence imaging of coronal section. Side images: magnified view of FUS targeted region (left) and contralateral site (right) in the middle image. Scale bars correspond to 500 μm (middle) and 100 μm (side), respectively.

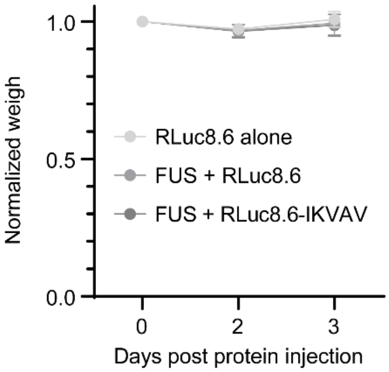

**Supplementary Figure 10. Mouse body weight analysis after systemic administration of unmodified or engineered RLuc8.6.** The weight just before intravenous injection of unmodified or engineered RLuc8.6 was used to normalize the weight of each mouse during the experiment. Data are presented as mean ± s.d. $n$ = 5 mice for each group. Intra-group differences between day 0 and 3 days after protein injection were identified using the Two-way ANOVA test. The statistical results were summarized in **Supplementary Table 2**.

**Table 1. The statistical results of intra-group differences between day 0 and 2 days after systemic administration of unmodified or engineered RLuc8.6 and AADC-IKVAV.**

| Group | Number of mice | Weight loss | P value | Significance |
|---|---|---|---|---|
| FUS alone | 11 | -2.3±1.7% | 0.0974 | ns |
| RLuc8.6 alone | 10 | 2.1±0.6% | 0.2148 | ns |
| FUS + RLuc8.6 | 10 | 3.4±0.7% | 0.0071 | ** |
| FUS + RLuc8.6-IKVAV | 10 | 3.2±0.8% | 0.0118 | * |
| AADC-IKVAV alone | 6 | 13.8±0.8% | <0.0001 | **** |
| FUS + AADC-IKVAV | 14 | 10.3±0.5% | <0.0001 | **** |

Statistical significance was denoted as $^*P < 0.05$, $^{**}P < 0.01$, $^{****}P < 0.0001$, and ns (not significant), using Two-way ANOVA test.

**Table 2. The statistical results of intra-group differences between day 0 and 3 days after systemic administration of unmodified or engineered RLuc8.6.**

| Group | Number of mice | P value | Significance |
|---|---|---|---|
| RLuc8.6 alone | 5 | 0.9043 | ns |
| FUS + RLuc8.6 | 5 | 0.9521 | ns |
| FUS + RLuc8.6-IKVAV | 5 | 0.6775 | ns |

Statistical significance was denoted as ns (not significant), using Two-way ANOVA test.

**Table 3. Synthetic oligos used for plasmid construction in this study.**

| | Vector backbone | Name | Sequences (5' to 3') | Restriction sites |
|---|---|---|---|---|
| RLuc8.6 | pRSETb | RLuc8.6 F | TGTACGACGATGACGATAA*GGATCC*GATGGCTT<br>CCAAGGTGTACGACCCCG | *BamHI* or *EcoRI* sites are underlined. |
| | | RLuc8.6 R | AGCAGCCGGATCAAGCTTC*GAATTC*TTACTGCT<br>CGTTCTTCAGCACGCGCT | |
| RLuc8.6-IKVAV | pRSETb | RLuc8.6 F | TGTACGACGATGACGATAA*GGATCC*GATGGCTT<br>CCAAGGTGTACGACCCCG | *BamHI* or *EcoRI* sites are underlined. |
| | | RLuc8.6 R | CTGCTCGTTCTTCAGCACGCGCT | |
| | | DNA fragment of IKVAV | GAGCGCGTGCTGAAGAACGAGCAGGGAGGAAG<br>TGGCAGCTCTGGCGGCAGTGGAGGGTCTGGTG<br>GCAGCGGAATCAAAGTCGCCGTGTAA<br>*GAATTC*GAAGCTTGATCCGGCTGCTAACAA | |
| RLuc8.6-YIGSR | pRSETb | RLuc8.6 F | TGTACGACGATGACGATAA*GGATCC*GATGGCTT<br>CCAAGGTGTACGACCCCG | *BamHI* or *EcoRI* sites are underlined. |
| | | RLuc8.6 R | CTGCTCGTTCTTCAGCACGCGCT | |
| | | DNA fragment of YIGSR | GAGCGCGTGCTGAAGAACGAGCAGGGAGGAAG<br>TGGCAGCTCTGGCGGCAGTGGAGGGTCTGGTG<br>GCAGCGGATACATTGGTTCCAGATAA<br>*GAATTC*GAAGCTTGATCCGGCTGCTAACAA | |
| RLuc8.6-GRGDS | pRSETb | RLuc8.6 F | TGTACGACGATGACGATAA*GGATCC*GATGGCTT<br>CCAAGGTGTACGACCCCG | *BamHI* or *EcoRI* sites are underlined. |
| | | RLuc8.6 R | CTGCTCGTTCTTCAGCACGCGCT | |
| | | DNA fragment of GRGDS | GAGCGCGTGCTGAAGAACGAGCAGGGAGGAAG<br>TGGCAGCTCTGGCGGCAGTGGAGGGTCTGGTG<br>GCAGCGGAGGTCGTGGTGATAGTTAA<br>*GAATTC*GAAGCTTGATCCGGCTGCTAACAA | |
| AADC-IKVAV | pRSETb | AADC-IKVAV F | TGTACGACGATGACGATAA*GGATCC*GATGAACG<br>CAAGTGAATTCCGAAGGAGAGGGA | *BamHI* or *EcoRI* sites are underlined. |
| | | AADC-IKVAV R | AGCAGCCGGATCAAGCTTC*GAATTC*TTACACGG<br>CGACTTTGATTCCGCTGCCA | |
| AADC-IKVAV template | pTrcHisA | AADC F | TGACGATAAGGATCGATGG*GGATCC*ATGAACGC<br>AAGTGAATTCCGAAGGAGAGGGA | *BamHI* or *KpnI* sites are underlined. |
| | | AADC R | CTCCCTCTCTGCTCGCAGCACG | |
| | | DNA fragment of IKVAV | ACGTGCTGCGAGCAGAGAGGGAGGGAGGAAGT<br>GGCAGCTCTGGCGGCAGTGGAGGGTCTGGTGG<br>CAGCGGAATCAAAGTCGCCGTGTAG*G*<br>*GTACC*ATATGGGAATTCGAAGCTTGGCTGT | |

Note: We first subcloned AADC-IKVAV into vector pTrcHisA, but only observed low protein production in *Escherichia coli*. However, it serves as a template for creating the expression plasmid of AADC-IKVAV using vector pRSETb, resulting in a better yield.

**Table 4. Protein coding sequences in this study.**

| Name | Protein coding sequence (CDS) or vector sequence | Note |
|---|---|---|
| RLuc8.6 | ATGCGGGGTTCTCATCATCATCATCATCATGGTATGGCTAGCATGACTGGTGGACAG<br>CAAATGGGTCGGGATCTGTACGACGATGACGATAAGGATCCGATGGCTTCCAAGGTG<br>TACGACCCCGAGCAACGCAAACGCATGATCACTGGGCCTCAGTGGTGGGCTCGCTG<br>CAAGCAAATGAACGTGCTGGACTCCTTCATCAACTACTATGATTCCGAGAAGCACGCC<br>GAGAACGCCGTGATTTTTCTGCATGGTAACGCTACCTCCAGCTACCTGTGGAGGCAC<br>GTCGTGCCTCACATCGAGCCCGTGGCTAGATGCATCATCCCTGATCTGATCGGAATG<br>GGTAAGTCCGGCAAGAGCGGGAATGGCTCATATCGCCTCCTGGATCACTACAAGTAC<br>CTCACCGCTTGGTTCGAGCTGCTGAACCTTCCAAAGAAAATCATCTTTGTGGGCCACG<br>ACTGGGGGAGCGCTCTGGCCTTTCACTACGCCTACGAGCACCAAGACAGGATCAAG<br>GCCATCGTCCATATGGAGAGTGTCGTGGACGTGATCGAGTCCTGGATGGGGTGGCC<br>TGACATCGAGGAGGAGCTGGCCCTGATCAAGAGCGAAGAGGGCGAGAAAATGGTGC<br>TTGAGAATAACTTCTTCGTCGAGACCCTGTTGCCAAGCAAGATCATGCGGAAACTGGA<br>GCCTGAGGAGTTCGCTGCCTACCTGGAGCCATTCAAGGAGAAGGGCGAGGTTAGAC<br>GGCCTACCCTCTCCTGGCCTCGCGAGATCCCTCTCGTTAAGGGAGGCAAGCCCGAC<br>GTCGTCCAGATTGTCCGCAACTACAACGCCTACCTTCGGGCCAGCGACGATCTGCCT<br>AAGCTGTTCATCGAGTCCGACCCTGGGTTCTTTTCCAACGCTATTGTCGAGGGAGCTA<br>AGAAGTTCCCTAACACCGAGTTCGTGAAGGTGAAGGGCCTCCACTTCCTCCAGGAGG<br>ACGCTCCAGATGAAATGGGTAAGTACATCAAGAGCTTCGTGGAGCGCGTGCTGAAGA<br>ACGAGCAG | His-tag sequence is highlighted in red, and RLuc8.6 in blue. Other sequences are start codon, T7 tag and X-press tag on the original vector pRSETb. |
| RLuc8.6-IKVAV | ATGCGGGGTTCTCATCATCATCATCATCATGGTATGGCTAGCATGACTGGTGGACAG<br>CAAATGGGTCGGGATCTGTACGACGATGACGATAAGGATCCGATGGCTTCCAAGGTG<br>TACGACCCCGAGCAACGCAAACGCATGATCACTGGGCCTCAGTGGTGGGCTCGCTG<br>CAAGCAAATGAACGTGCTGGACTCCTTCATCAACTACTATGATTCCGAGAAGCACGCC<br>GAGAACGCCGTGATTTTTCTGCATGGTAACGCTACCTCCAGCTACCTGTGGAGGCAC<br>GTCGTGCCTCACATCGAGCCCGTGGCTAGATGCATCATCCCTGATCTGATCGGAATG<br>GGTAAGTCCGGCAAGAGCGGGAATGGCTCATATCGCCTCCTGGATCACTACAAGTAC<br>CTCACCGCTTGGTTCGAGCTGCTGAACCTTCCAAAGAAAATCATCTTTGTGGGCCACG<br>ACTGGGGGAGCGCTCTGGCCTTTCACTACGCCTACGAGCACCAAGACAGGATCAAG<br>GCCATCGTCCATATGGAGAGTGTCGTGGACGTGATCGAGTCCTGGATGGGGTGGCC<br>TGACATCGAGGAGGAGCTGGCCCTGATCAAGAGCGAAGAGGGCGAGAAAATGGTGC<br>TTGAGAATAACTTCTTCGTCGAGACCCTGTTGCCAAGCAAGATCATGCGGAAACTGGA<br>GCCTGAGGAGTTCGCTGCCTACCTGGAGCCATTCAAGGAGAAGGGCGAGGTTAGAC<br>GGCCTACCCTCTCCTGGCCTCGCGAGATCCCTCTCGTTAAGGGAGGCAAGCCCGAC<br>GTCGTCCAGATTGTCCGCAACTACAACGCCTACCTTCGGGCCAGCGACGATCTGCCT<br>AAGCTGTTCATCGAGTCCGACCCTGGGTTCTTTTCCAACGCTATTGTCGAGGGAGCTA<br>AGAAGTTCCCTAACACCGAGTTCGTGAAGGTGAAGGGCCTCCACTTCCTCCAGGAGG<br>ACGCTCCAGATGAAATGGGTAAGTACATCAAGAGCTTCGTGGAGCGCGTGCTGAAGA<br>ACGAGCAGGGAGGAAGTGGCAGCTCTGGCGGCAGTGGAGGGTCTGGTGGCAGCGG<br>AATCAAAGTCGCCGTG | His-tag sequence is highlighted in red, RLuc8.6 in blue, GS linker in green, and IKVAV in purple. Other sequences are start codon, T7 tag and X-press tag on the original vector pRSETb. |
| RLuc8.6-YIGSR | ATGCGGGGTTCTCATCATCATCATCATCATGGTATGGCTAGCATGACTGGTGGACAG<br>CAAATGGGTCGGGATCTGTACGACGATGACGATAAGGATCCGATGGCTTCCAAGGTG<br>TACGACCCCGAGCAACGCAAACGCATGATCACTGGGCCTCAGTGGTGGGCTCGCTG<br>CAAGCAAATGAACGTGCTGGACTCCTTCATCAACTACTATGATTCCGAGAAGCACGCC<br>GAGAACGCCGTGATTTTTCTGCATGGTAACGCTACCTCCAGCTACCTGTGGAGGCAC<br>GTCGTGCCTCACATCGAGCCCGTGGCTAGATGCATCATCCCTGATCTGATCGGAATG<br>GGTAAGTCCGGCAAGAGCGGGAATGGCTCATATCGCCTCCTGGATCACTACAAGTAC<br>CTCACCGCTTGGTTCGAGCTGCTGAACCTTCCAAAGAAAATCATCTTTGTGGGCCACG<br>ACTGGGGGAGCGCTCTGGCCTTTCACTACGCCTACGAGCACCAAGACAGGATCAAG<br>GCCATCGTCCATATGGAGAGTGTCGTGGACGTGATCGAGTCCTGGATGGGGTGGCC<br>TGACATCGAGGAGGAGCTGGCCCTGATCAAGAGCGAAGAGGGCGAGAAAATGGTGC<br>TTGAGAATAACTTCTTCGTCGAGACCCTGTTGCCAAGCAAGATCATGCGGAAACTGGA<br>GCCTGAGGAGTTCGCTGCCTACCTGGAGCCATTCAAGGAGAAGGGCGAGGTTAGAC<br>GGCCTACCCTCTCCTGGCCTCGCGAGATCCCTCTCGTTAAGGGAGGCAAGCCCGAC<br>GTCGTCCAGATTGTCCGCAACTACAACGCCTACCTTCGGGCCAGCGACGATCTGCCT<br>AAGCTGTTCATCGAGTCCGACCCTGGGTTCTTTTCCAACGCTATTGTCGAGGGAGCTA<br>AGAAGTTCCCTAACACCGAGTTCGTGAAGGTGAAGGGCCTCCACTTCCTCCAGGAGG<br>ACGCTCCAGATGAAATGGGTAAGTACATCAAGAGCTTCGTGGAGCGCGTGCTGAAGA<br>ACGAGCAGGGAGGAAGTGGCAGCTCTGGCGGCAGTGGAGGGTCTGGTGGCAGCGG<br>ATACATTGGTTCCAGA | His-tag sequence is highlighted in red, RLuc8.6 in blue, GS linker in green, and YIGSR in purple. Other sequences are start codon, T7 tag and X-press tag on the original vector pRSETb. |
| RLuc8.6-GRGDS | ATGCGGGGTTCTCATCATCATCATCATCATGGTATGGCTAGCATGACTGGTGGACAG<br>CAAATGGGTCGGGATCTGTACGACGATGACGATAAGGATCCGATGGCTTCCAAGGTG<br>TACGACCCCGAGCAACGCAAACGCATGATCACTGGGCCTCAGTGGTGGGCTCGCTG<br>CAAGCAAATGAACGTGCTGGACTCCTTCATCAACTACTATGATTCCGAGAAGCACGCC<br>GAGAACGCCGTGATTTTTCTGCATGGTAACGCTACCTCCAGCTACCTGTGGAGGCAC<br>GTCGTGCCTCACATCGAGCCCGTGGCTAGATGCATCATCCCTGATCTGATCGGAATG<br>GGTAAGTCCGGCAAGAGCGGGAATGGCTCATATCGCCTCCTGGATCACTACAAGTAC | His-tag sequence is highlighted in red, RLuc8.6 in blue, GS linker in green, and GRGDS in purple. Other sequences are start codon, T7 tag |

| | | |
|---|---|---|
| | CTCACCGCTTGGTTCGAGCTGCTGAACCTTCCAAAGAAAATCATCTTTGTGGGCCACG<br>ACTGGGGGAGCGCTCTGGCCTTTCACTACGCCTACGAGCACCAAGACAGGATCAAG<br>GCCATCGTCCATATGGAGAGTGTCGTGGACGTGATCGAGTCCTGGATGGGGTGGCC<br>TGACATCGAGGAGGAGCTGGCCCTGATCAAGAGCGAAGAGGGCGAGAAAATGGTGC<br>TTGAGAATAACTTCTTCGTCGAGACCCTGTTGCCAAGCAAGATCATGCGGAAACTGGA<br>GCCTGAGGAGTTCGCTGCCTACCTGGAGCCATTCAAGGAGAAGGGCGAGGTTAGAC<br>GGCCTACCCTCTCCTGGCCTCGCGAGATCCCTCTCGTTAAGGGAGGCAAGCCCGAC<br>GTCGTCCAGATTGTCCGCAACTACAACGCCTACCTTCGGGCCAGCGACGATCTGCCT<br>AAGCTGTTCATCGAGTCCGACCCTGGGTTCTTTTCCAACGCTATTGTCGAGGGAGCTA<br>AGAAGTTCCCTAACACCGAGTTCGTGAAGGTGAAGGGCCTCCACTTCCTCCAGGAGG<br>ACGCTCCAGATGAAATGGGTAAGTACATCAAGAGCTTCGTGGAGCGCGTGCTGAAGA<br>ACGAGCAGGGAGGAAGTGGCAGCTCTGGCGGCAGTGGAGGGTCTGGTGGCAGCGG<br>AGGTCGTGGTGATAGT | and X-press tag on the original vector pRSETb. |
| AADC-IKVAV | ATGCGGGGTTCTCATCATCATCATCATCATGGTATGGCTAGCATGACTGGTGGACAG<br>CAAATGGGTCGGGATCTGTACGACGATGACGATAAGGATCCGATGAACGCAAGTGAA<br>TTCCGAAGGAGAGGGAAGGAGATGGTGGATTACATGGCCAACTACATGGAAGGCATT<br>GAGGGACGCCAGGTCTACCCTGACGTGGAGCCCGGGTACCTGCGGCCGCTGATCCC<br>TGCCGCTGCCCCTCAGGAGCCAGACACGTTTGAGGACATCATCAACGACGTTGAGAA<br>GATAATCATGCCTGGGGTGACGCACTGGCACAGCCCCTACTTCTTCGCCTACTTCCC<br>CACTGCCAGCTCGTACCCGGCCATGCTTGCGGACATGCTGTGCGGGGCCATTGGCT<br>GCATCGGCTTCTCCTGGGCGGCAAGCCCAGCATGCACAGAGCTGGAGACTGTGATG<br>ATGGACTGGCTCGGGAAGATGCTGGAACTACCAAAGGCATTTTTGAATGAGAAAGCT<br>GGAGAAGGGGGAGGAGTGATCCAGGGAAGTGCCAGTGAAGCCACCCTGGTGGCCC<br>TGCTGGCCGCTCGGACCAAAGTGATCCATCGGCTGCAGGCAGCGTCCCCAGAGCTC<br>ACACAGGCCGCTATCATGGAGAAGCTGGTGGCTTACTCATCCGATCAGGCACACTCC<br>TCAGTGGAAAGAGCTGGGTTAATTGGTGGAGTGAAATTAAAAGCCATCCCCTCAGAT<br>GGCAACTTCGCCATGCGTGCGTCTGCCCTGCAGGAAGCCCTGGAGAGAGACAAAGC<br>GGCTGGCCTGATTCCTTTCTTTATGGTTGCCACCCTGGGGACCACAACATGCTGCTC<br>CTTTGACAATCTCTTAGAAGTCGGTCCTATCTGCAACAAGGAAGACATATGGCTGCAC<br>GTTGATGCAGCCTACGCAGGCAGTGCATTCATCTGCCCTGAGTTCCGGCACCTTCTG<br>AATGGAGTGGAGTTTGCAGATTCATTCAACTTTAATCCCCACAAATGGCTATTGGTGA<br>ATTTTGACTGTTCTGCCATGTGGGTGAAAAAGAGAACAGACTTAACGGGAGCCTTTAG<br>ACTGGACCCCACTTACCTGAAGCACAGCCATCAGGATTCAGGGCTTATCACTGACTA<br>CCGGCATTGGCAGATACCACTGGGCAGAAGATTTCGCTCTTTGAAAATGTGGTTTGTA<br>TTTAGGATGTATGGAGTCAAAGGACTGCAGGCTTATATCCGCAAGCATGTCCAGCTGT<br>CCCATGAGTTTGAGTCACTGGTGCGCCAGGATCCCCGCTTTGAAATCTGTGTGGAAG<br>TCATTCTGGGGCTTGTCTGCTTTCGGCTAAAGGGTTCCAACAAAGTGAATGAAGCTCT<br>TCTGCAAAGAATAAACAGTGCCAAAAAAATCCACTTGGTTCCATGTCACCTCAGGGAC<br>AAGTTTGTCCTGCGCTTTGCCATCTGTTCTCGCACGGTGGAATCTGCCCATGTGCAG<br>CGGGCCTGGGAACACATCAAAGAGCTGGCGGCCGACGTGCTGCGAGCAGAGAGGG<br>AGGGAGGAAGTGGCAGCTCTGGCGGCAGTGGAGGGTCTGGTGGCAGCGGAATCAA<br>AGTCGCCGTG | His-tag sequence is highlighted in red, AADC in blue, GS linker in green, and IKVAV in purple. Other sequences are start codon, T7 tag and X-press tag on the original vector pRSETb. |
| pRSETb vector | TTCATTTTTAATTTAAAAGGATCTAGGTGAAGATCCTTTTTGATAATCTCATGACCAAAA<br>TCCCTTAACGTGAGTTTTCGTTCCACTGAGCGTCAGACCCCGTAGAAAAGATCAAAGG<br>ATCTTCTTGAGATCCTTTTTTTCTGCGCGTAATCTGCTGCTTGCAAACAAAAAAACCAC<br>CGCTACCAGCGGTGGTTTGTTTGCCGGATCAAGAGCTACCAACTCTTTTTCCGAAGGT<br>AACTGGCTTCAGCAGAGCGCAGATACCAAATACTGTTCTTCTAGTGTAGCCGTAGTTA<br>GGCCACCACTTCAAGAACTCTGTAGCACCGCCTACATACCTCGCTCTGCTAATCCTGT<br>TACCAGTGGCTGCTGCCAGTGGCGATAAGTCGTGTCTTACCGGGTTGGACTCAAGAC<br>GATAGTTACCGGATAAGGCGCAGCGGTCGGGCTGAACGGGGGGTTCGTGCACACAG<br>CCCAGCTTGGAGCGAACGACCTACACCGAACTGAGATACCTACAGCGTGAGCTATGA<br>GAAAGCGCCACGCTTCCCGAAGGGAGAAAGGCGGACAGGTATCCGGTAAGCGGCAG<br>GGTCGGAACAGGAGAGCGCACGAGGGAGCTTCCAGGGGGAAACGCCTGGTATCTTT<br>ATAGTCCTGTCGGGTTTCGCCACCTCTGACTTGAGCGTCGATTTTTGTGATGCTCGTC<br>AGGGGGGCGGAGCCTATGGAAAAACGCCAGCAACGCGGCCTTTTTACGGTTCCTGG<br>CCTTTTGCTGGCCTTTTGCTCACATGTTCTTTCCTGCGTTATCCCCTGATTCTGTGGAT<br>AACCGTATTACCGCCTTTGAGTGAGCTGATACCGCTCGCCGCAGCCGAACGACCGAG<br>CGCAGCGAGTCAGTGAGCGAGGAAGCGGAAGAGCGCCCAATACGCAAACCGCCTCT<br>CCCCGCGCGTTGGCCGATTCATTAATGCAGGATCTCGATCCCGCGAAATTAATACGA<br>CTCACTATAGGGAGACCACAACGGTTTCCCTCTAGAAATAATTTTGTTTAACTTTAAGA<br>AGGAGATATACATATGCGGGGTTCTCATCATCATCATCATCATGGTATGGCTAGCATG<br>ACTGGTGGACAGCAAATGGGTCGGGATCTGTACGACGATGACGATAAGGATCCGATG<br>GTGAGCAAGGGCGAGGAGGTCATCAAAGAGTTCATGCGCTTCAAGGTGCGCATGGA<br>GGGCTCCATGAACGGCCACGAGTTCGAGATCGAGGGCGAGGGCGAGGGCCGCCCC<br>TACGAGGGCACCCAGACCGCCAAGCTGAAGGTGACCAAGGGCGGCCCCCTGCCCTT<br>CGCCTGGGACATCCTGTCCCCCCAGTTCATGTACGGCTCCAAGGCGTACGTGAAGCA<br>CCCCGCCGACATCCCCGATTACAAGAAGCTGTCCTTCCCCGAGGGCTTCAAGTGGGA<br>GCGCGTGATGAACTTCGAGGACGGCGGTCTGGTGACCGTGACCCAGGACTCCTCCC<br>TGCAGGACGGCACGCTGATCTACAAGGTGAAGATGCGCGGCACCAACTTCCCCCCC<br>GACGGCCCCGTAATGCAGAAGAAGACCATGGGCTGGGAGGCCTCCACCGAGCGCCT | |

| | |
|---|---|
| | GTACCCCCGCGACGGCGTGCTGAAGGGCGAGATCCACCAGGCCCTGAAGCTGAAGG<br>ACGGCGGCCACTACCTGGTGGAGTTCAAGACCATCTACATGGCCAAGAAGCCCGTG<br>CAACTGCCCGGCTACTACTACGTGGACACCAAGCTGGACATCACCTCCCACAACGAG<br>GACTACACCATCGTGGAACAGTACGAGCGCTCCGAGGGCCGCCACCACCTGTTCCT<br>GGGGCATGGCACCGGCAGCACCGGCAGCGGCAGCTCCGGCACCGCCTCCTCCGAG<br>GACAACAACATGGCCGTCATCAAAGAGTTCATGCGCTTCAAGGTGCGCATGGAGGGC<br>TCCATGAACGGCCACGAGTTCGAGATCGAGGGCGAGGGCGAGGGCCGCCCCTACG<br>AGGGCACCCAGACCGCCAAGCTGAAGGTGACCAAGGGCGGCCCCCTGCCCTTCGCC<br>TGGGACATCCTGTCCCCCCAGTTCATGTACGGCTCCAAGGCGTACGTGAAGCACCCC<br>GCCGACATCCCCGATTACAAGAAGCTGTCCTTCCCCGAGGGCTTCAAGTGGGAGCG<br>CGTGATGAACTTCGAGGACGGCGGTCTGGTGACCGTGACCCAGGACTCCTCCCTGC<br>AGGACGGCACGCTGATCTACAAGGTGAAGATGCGCGGCACCAACTTCCCCCCCGAC<br>GGCCCCGTAATGCAGAAGAAGACCATGGGCTGGGAGGCCTCCACCGAGCGCCTGTA<br>CCCCCGCGACGGCGTGCTGAAGGGCGAGATCCACCAGGCCCTGAAGCTGAAGGAC<br>GGCGGCCACTACCTGGTGGAGTTCAAGACCATCTACATGGCCAAGAAGCCCGTGCAA<br>CTGCCCGGCTACTACTACGTGGACACCAAGCTGGACATCACCTCCCACAACGAGGAC<br>TACACCATCGTGGAACAGTACGAGCGCTCCGAGGGCCGCCACCACCTGTTCCGGCT<br>GGAAGATTTCGTTGGGGACTGGCGACAGACAGCCGGCTACAACCTGGACCAAGTCC<br>TTGAACAGGGAGGTGTGTCCAGTTTGTTTCAGAATCTCGGGGTGTCCGTAACTCCGA<br>TCCAAAGGATTGTCCTGAGCGGTGAAAATGGGCTGAAGATCGACATCCATGTCATCAT<br>CCCGTATGAAGGTCTGAGCGGCGACCAAATGGGCCAGATCGAAAAAATTTTTAAGGT<br>GGTGTACCCTGTGGATGATCATCACTTTAAGGTGATCCTGCACTATGGCACACTGGTA<br>ATCGACGGGGTTACGCCGAACATGATCGACTATTTCGGACGGCCGTATGAAGGCATC<br>GCCGTGTTCGACGGCAAAAAGATCACTGTAACAGGGACCCTGTGGAACGGCAACAAA<br>ATTATCGACGAGCGCCTGATCAACCCCGACGGCTCCCTGCTGTTCCGAGTAACCATC<br>AACGGAGTGACCGGCTGGCGGCTGTGCGAACGCATTCTGGCGTAAGAATTCGAAGC<br>TTGATCCGGCTGCTAACAAAGCCCGAAAGGAAGCTGAGTTGGCTGCTGCCACCGCTG<br>AGCAATAACTAGCATAACCCCTTGGGGCCTCTAAACGGGTCTTGAGGGGTTTTTTGCT<br>GAAAGGAGGAACTATATCCGGATCTGGCGTAATAGCGAAGAGGCCCGCACCGATCG<br>CCCTTCCCAACAGTTGCGCAGCCTGAATGGCGAATGGGACGCGCCCTGTAGCGGCG<br>CATTAAGCGCGGCGGGTGTGGTGGTTACGCGCAGCGTGACCGCTACACTTGCCAGC<br>GCCCTAGCGCCCGCTCCTTTCGCTTTCTTCCCTTCCTTTCTCGCCACGTTCGCCGGCT<br>TTCCCCGTCAAGCTCTAAATCGGGGGCTCCCTTTAGGGTTCCGATTTAGTGCTTTACG<br>GCACCTCGACCCCAAAAAACTTGATTAGGGTGATGGTTCACGTAGTGGGCCATCGCC<br>CTGATAGACGGTTTTTCGCCCTTTGACGTTGGAGTCCACGTTCTTTAATAGTGGACTC<br>TTGTTCCAAACTGGAACAACACTCAACCCTATCTCGGTCTATTCTTTTGATTTATAAGG<br>GATTTTGCCGATTTCGGCCTATTGGTTAAAAAATGAGCTGATTTAACAAAAATTTAACG<br>CGAATTTTAACAAAATATTAACGCTTACAATTTAGGTGGCACTTTTCGGGGAAATGTGC<br>GCGGAACCCCTATTTGTTTATTTTTCTAAATACATTCAAATATGTATCCGCTCATGAGA<br>CAATAACCCTGATAAATGCTTCAATAATATTGAAAAAGGAAGAGTATGAGTATTCAACA<br>TTTCCGTGTCGCCCTTATTCCCTTTTTTGCGGCATTTTGCCTTCCTGTTTTTGCTCACC<br>CAGAAACGCTGGTGAAAGTAAAAGATGCTGAAGATCAGTTGGGTGCACGAGTGGGTT<br>ACATCGAACTGGATCTCAACAGCGGTAAGATCCTTGAGAGTTTTCGCCCCGAAGAAC<br>GTTTTCCAATGATGAGCACTTTTAAAGTTCTGCTATGTGGCGCGGTATTATCCCGTATT<br>GACGCCGGGCAAGAGCAACTCGGTCGCCGCATACACTATTCTCAGAATGACTTGGTT<br>GAGTACTCACCAGTCACAGAAAAGCATCTTACGGATGGCATGACAGTAAGAGAATTAT<br>GCAGTGCTGCCATAACCATGAGTGATAACACTGCGGCCAACTTACTTCTGACAACGAT<br>CGGAGGACCGAAGGAGCTAACCGCTTTTTTGCACAACATGGGGGATCATGTAACTCG<br>CCTTGATCGTTGGGAACCGGAGCTGAATGAAGCCATACCAAACGACGAGCGTGACAC<br>CACGATGCCTGTAGCAATGGCAACAACGTTGCGCAAACTATTAACTGGCGAACTACTT<br>ACTCTAGCTTCCCGGCAACAATTAATAGACTGGATGGAGGCGGATAAAGTTGCAGGA<br>CCACTTCTGCGCTCGGCCCTTCCGGCTGGCTGGTTTATTGCTGATAAATCTGGAGCC<br>GGTGAGCGTGGGTCTCGCGGTATCATTGCAGCACTGGGGCCAGATGGTAAGCCCTC<br>CCGTATCGTAGTTATCTACACGACGGGGAGTCAGGCAACTATGGATGAACGAAATAG<br>ACAGATCGCTGAGATAGGTGCCTCACTGATTAAGCATTGGTAACTGTCAGACCAAGTT<br>TACTCATATATACTTTAGATTGATTTAAAAC |

**Table 5. FUS target coordinates used for this study.**

| Related Figure | Experiment | FUS target | Medio-lateral (mm) | Anterior–posterior (mm) | Dorso-ventral (mm) |
|---|---|---|---|---|---|
| Fig. 2b-c | IHC analysis of RLuc8.6 (4 sites) | Site-1 | -2.74 | -0.1 | 4.33 |
| | | Site-2 | -1.85 | -0.1 | 4 |
| | | Site-3 | -2.5 | 0.13 | 4.48 |
| | | Site-4 | -1.6 | 0.13 | 3.95 |
| Fig. 2d-e | *In vivo* BLI (1 site) | Site-1 | -2.42 | -0.12 | 4.3 |
| Fig. 3 | *Ex vivo* BLI (3 sites) | Site-1 | -2.42 | -0.45 | 3.43 |
| | | Site-2 | -1.3 | -1.57 | 2.88 |
| | | Site-3 | -1.22 | -3.8 | 2.88 |
| Fig. 4 | c-Fos activation (1 site) | Site-1 | -2.42 | -0.12 | 4.1 |
| Fig. 5 | Behavior test (2 sites) | Site-1 | -2.32 | 0.18 | 4 |
| | | Site-2 | -2.92 | -0.62 | 4 |

**Movies 1-2: Representative videos of open field test for a mouse treated with FUS + AADC-IKVAV before (Move 1) and after (Move 2) administration of L-DOPA.**
